# A community benchmark enables high-resolution, cross-lab quantification of mouse social behavior

**DOI:** 10.64898/2026.10.18.758876

**Authors:** Markus Marks, Ruize Yang, Ziang Liu, Ryan Ye, Lara K. Scaria, Sam A. Golden, Jennifer J. Sun, Talmo Pereira, Ann Kennedy, Days of our Mice Consortium

## Abstract

Social behavior is a sensitive readout of brain state, but quantifying rodent social behavior at scale has relied on siloed, lab-specific tools. We assembled Days of our Mice (DooM), a benchmark of manually annotated video and pose tracking spanning 15 laboratories and 33 social and solitary behaviors, and hosted a community behavior classification challenge. We built on challenge solutions to develop the High-Dimensional Rodent Annotator (HiDRA), a behavior classification tool that generalizes to labs unseen during training and learns cross-lab movement features. Applied without retraining to experimental data, HiDRA resolves line-specific and state-dependent behavioral fingerprints with strong generalizability and reproducibility, establishing high-resolution, shareable social phenotyping as a scalable readout of brain and internal state. We release the DooM dataset and HiDRA classifiers as open resources.

## Introduction

Control of social behavior is an essential brain function that is inextricably linked to survival. It engages systems for sensory processing, learning and memory, planning and cognition, context-dependent decision-making, and motor control, and it shifts with an animal’s genetic background, development, experience, and internal state(*1–4*). Social behavior thus holds a privileged position as a highly attuned lens into brain state(*5–7*): its alteration is an indicator of dysfunction across the lifespan, for example in autism, schizophrenia, mood disorders, and neurodegeneration(*8–14*), and of state perturbations such as pain, stress, hunger, and illness(*15–18*). Mouse models remain an essential means of linking these effects to underlying biological mechanisms.

Socially interacting animals produce a rich and time-evolving repertoire of actions. Standard social behavior assays like the three-chamber test sacrifice this richness for ease of throughput(*19*, *20*). The alternative approach of studying free social interactions historically relied on manual, frame-by-frame scoring of animals’ actions, but has been supplemented in recent years by machine learning(*21–31*). In supervised behavior classification, human-defined behaviors are detected from features of animal pose(*32*, *33*) or video; supervised classifiers can match human performance given adequate training examples(*24*, *34*). But supervised methods rely on costly human-generated training data and do not generalize to new experimental settings(*35*, *36*). Thus, while machine learning promises to accelerate behavior analysis, trained models for classification and the definitions of actions themselves still remain siloed within labs(*30*).

Whether a common vocabulary of social behavior can be detected and quantified across diverse experimental settings is unknown. To answer this question, we assembled Days of Our Mice (DooM), a dataset of manually annotated video and pose tracking contributed by 15 laboratories, spanning 180 hours and 33 social and solitary behaviors, and challenged the machine learning community to build classifiers from it. Building on the winning approach, we trained the High-Dimensional Rodent Annotator (HiDRA), a single model that scores all 33 behaviors according to each lab’s working definitions, producing 82 distinct classifier heads. HiDRA learns a shared representation of movement, from which the same behavior can be read out regardless of experimental setting. As a result, its classifiers can be transferred to new settings not seen during training, and are robust to variation in video resolution, frame rate, and appearance, outperforming direct video-based methods. Applied without retraining to test cases of social phenotyping in mouse lines with autism-associated mutations and in a multi-day experience assay, HiDRA reproduced and extended previously reported findings. We release DooM and HiDRA as open resources for high-resolution, shareable social phenotyping and as the new standard for pre-clinical social behavior benchmarking.

### The Days Of Our Mice (DooM) is a cross-laboratory dataset of mouse social behavior

Previous tools and benchmarks(*37–42*) for rodent behavior quantification have focused on single-lab settings; such methods fail on datasets that are out of the training distribution of one lab’s data, making it infeasible to share or combine classifiers across datasets. To overcome this limitation, we constructed the Days Of Our Mice (DooM), a multi-lab benchmark dataset for framewise action annotation in socially interacting mice (**Fig 1A**). DooM consists of over 600 top-view videos of mouse social interactions in diverse experimental settings (**Fig 1B**), accompanied by metadata, markerless pose estimation, and frame-by-frame manual behavior annotation.

**Figure 1.**
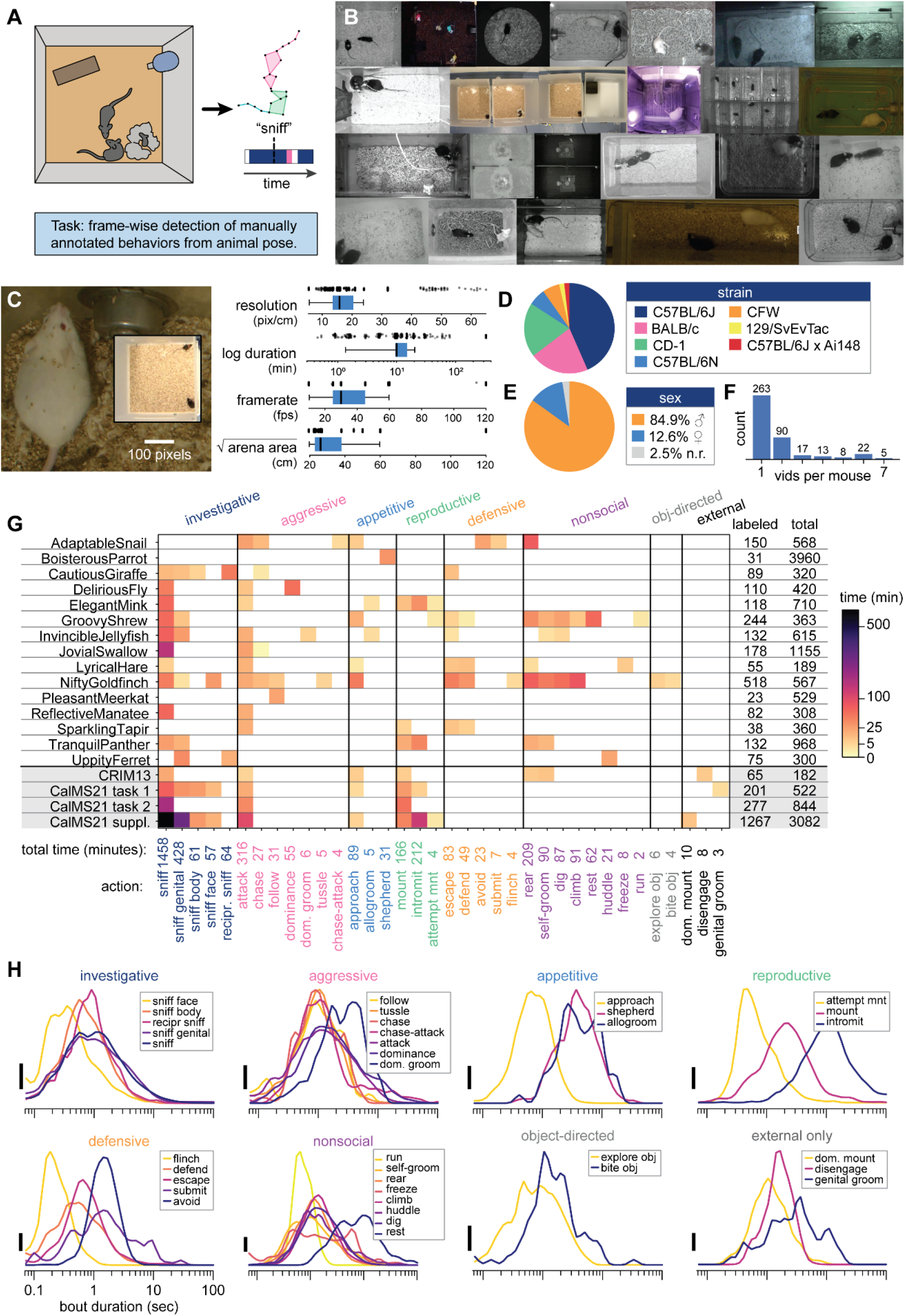
The Days Of Our Mice (DooM) benchmark dataset captures the diversity of mouse social behavior experiments. **A)** DooM is a dataset of videos, markerless pose estimates, and framewise manual annotations of behavior, to be used to benchmark models for supervised action detection in socially interacting mice. **B)** Representative video frames from the 15 contributor labs in the DooM dataset. The diversity of video appearances poses a challenge for automated action recognition that generalizes across labs. **C)** Left: Example images contrasting highest- and lowest-resolution videos in the dataset. Right: Distribution of video acquisition settings across the dataset; points are individual videos, box plots reflect median and interquartile range. **D)** Strain breakdown of mice in the dataset. **E)** Sex balance of mice in the dataset. **F)** Frequency of animal re-appearance across videos, in instances where individual identity was recorded; note that this reflects only a subset of videos in the dataset. **G)** Matrix of behaviors annotated per lab, colored by number of minutes of behavior annotated (square root color scale). Non-competition datasets are shown at the bottom of the table and marked in gray. Rightmost columns give the total number of annotated minutes from each lab and total duration of behavior videos. Note that on some video frames, none of the annotated behaviors are occurring. **H)** Distribution of bout durations for each behavior in the dataset, merged across labs and including the supplementary datasets (CRIM13 and CalMS21). Vertical scale bar is 2% probability density.

Our goal in constructing DooM was to capture the true heterogeneity of mouse social behavior experiments, and the dataset exhibits at least an order of magnitude variation in most dimensions of experimental metadata (**Fig 1C**). Biological variability across the dataset includes age, sex, strain, and experimental background of the animals (**Fig 1D-E**). While not all labs reported consistent cross-session identity for all animals, this data is also provided where known; a subset of animals appear across multiple assays, allowing for study of reidentifiability (**Fig 1F**). Each contributor lab maintains its own mouse pose representation (**Suppl Fig 1**), but we applied modest renaming of body parts for ease of use.

The videos in DooM are all derived from prior experiments, and the behaviors annotated reflect the choices and operational behavior definitions of those original studies. Labs used a total of 121 unique strings to identify behaviors across the dataset; after grammatical standardization and merging based on documentation and visual inspection, we condensed these to 46 unique behaviors, of which we identified a final subset of 33 that were annotated consistently and with sufficient examples for downstream analysis (see Methods). Of the 33 behaviors in DooM, 13 are unique to individual labs. The remainder were all annotated by at least two labs, permitting analysis of cross-lab agreement in behavior definitions. In total, the dataset contains 82 distinct lab-behavior pairs (**Fig 1G**). Annotated actions span a range of timescales, with typical durations on the order of milliseconds to seconds (**Fig 1H**).

### A community-scale benchmark for automated social behavior recognition

To define state-of-the-art performance for frame-wise behavior classification, we ran a two-month challenge on Kaggle using the DooM dataset (Methods, **Suppl Fig 2A**). We augmented available classifier training data by converting the previously published CalMS21(*38*), MABe22(*40*), and CRIM13(*37*) datasets into the DooM format, totaling 395 hours of data across 9,039 videos. We also updated CalMS21 with expanded manual annotations of behaviors for the Supplemental and Task 1 datasets.

The challenge drew over 30,000 submissions and 200 tutorial notebooks from 1,781 participants in 85 countries. The five top-ranked submissions achieved an average F1 score of 0.537 on the held-out test set (**Suppl Fig 2B**). The second-through fifth-place solutions used a combination of neural network-based approaches (recurrent neural networks (RNNs), graph neural networks (GNNs), and Transformers) and classical machine learning methods (gradient-boosted trees, via XGBoost), operating on hand-crafted features such as per-mouse and pairwise velocities, distances, and angles. The top submission stood out for its use of self-supervised learning for feature extraction, via a pair of GNNs trained to forecast keypoint movements for single animals and animal pairs. This advantage of learned over hand-crafted features is consistent with previous work(*31*, *43–45*).

### The High-Dimensional Rodent Annotator (HiDRA) captures the behavioral repertoire of freely interacting mice

Building on the competition outcome, we developed a lightly modified version of the first-place model, and retrained it following additional refinement of the DooM dataset (Methods). We call this retrained, 82-head model the High-Dimensional Rodent Annotator (HiDRA, **Fig 2A**). We used HiDRA to generate predicted ethograms of all behaviors in the DooM test set (**Fig 2B**), and evaluated their ability to recover common summary statistics used in behavioral analysis (**Fig 2C-D**). HiDRA achieves a per-lab macro-average F1 score of 0.66 when evaluated on the DooM test set (**Fig 2E**).

**Figure 2.**
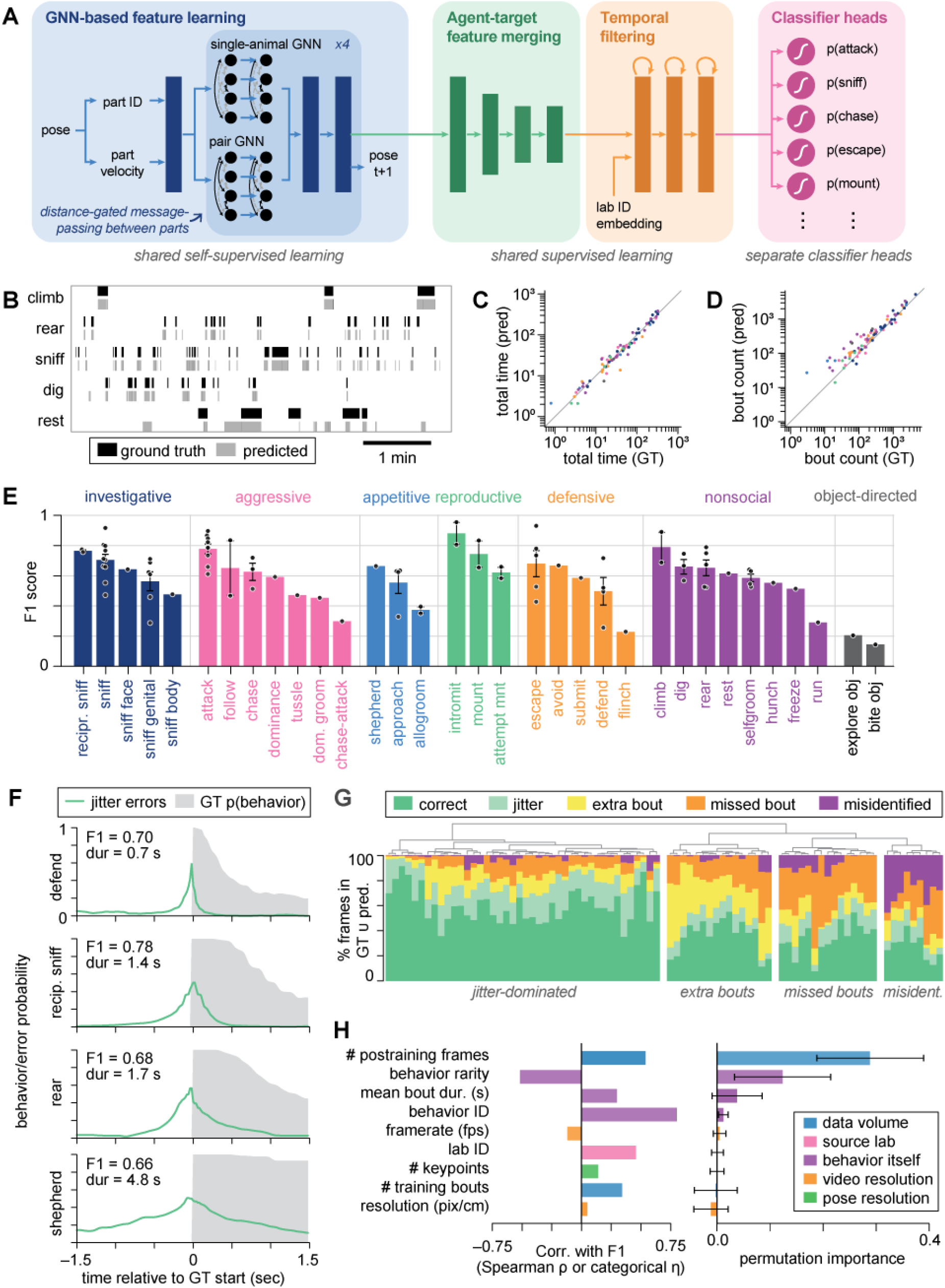
Supervised classifiers trained on self-supervised pose features detect a variety of mouse behaviors. **A)** HiDRA network architecture. **B)** Ethogram for a representative video showing human-annotated bouts (black) vs classifier predictions (gray) for five behaviors. **C)** HiDRA classifiers accurately capture time spent engaged in behavior, falling within a 10% margin of error in 29 of 82 classifiers. ( **D)** HiDRA classifiers modestly overestimate bout counts compared to humans. 12 of 82 classifiers fall within a 10% margin of error; merging bouts less than 500 ms apart increases this to 22 of 82. **E)** Classifier F1 score for the 33 behaviors in the DooM dataset. Points reflect classifiers from different labs trained for the same behavior. **F)** Classifier-detected behavior onset relative to human annotations, showing behavior-specific jitter in onset detection. Jitter levels are largely correlated with bout duration and speed of behavior. **G)** Breakdown of classifier error types contributing to F1 score, revealing heterogeneous basis of error across classifiers. High-performing classifiers are dominated by jitter errors (TP/FP frames occurring within 0.5 seconds of a GT bout). Errors related to extra bouts are candidates for human annotation error. **H)** Identification of metadata features that predict classifier F1 score. Left: Spearman’s ρ between metadata values and F1 score for each feature, sorted by magnitude. Right: permutation importance of metadata features in a cross-validated regression model trained to predict F1 score, showing training set size and bout duration to be the clearest drivers of classifier performance.

To ask whether reliance on 2d keypoints imposes a significant limit on classifier performance, we replaced HiDRA’s GNN feature extractor with features extracted directly from videos using the VideoPrism foundation model(*46*, *47*), and retrained the HiDRA supervised classification model on this rich feature input. Surprisingly, the pose-based model outperformed video-derived features for 80 out of 82 tested behaviors (**Suppl Fig 3**), indicating that pose keypoints are a highly effective representation for classification of the behaviors annotated in DooM. Future efforts to either improve video foundation models or fine-tune video foundation model backbones to mouse experimental data(*48–50*) may overcome these limitations; the DooM benchmark videos will be a useful resource for this effort.

### Identifying predictors of classifier performance across 82 lab-behavior pairs

The top-performing architectures in our competition had highly correlated F1 scores over behaviors (**Suppl Fig 2D**), suggesting that classification difficulty is a property of the data itself rather than of any particular architecture. We therefore used DooM and HiDRA to study at scale the factors that limit behavior classifier performance.

We hypothesized that one limiting factor in HiDRA classifier F1 score was annotation jitter: bout-to-bout variation in the timing of manual annotation onset, which arises because humans struggle to label transitions consistently(*24*, *51*, *52*). We decomposed the errors of all 82 HiDRA classifiers into extra bouts, missed bouts, misidentified behaviors, and jitter errors falling within half a second of a ground-truth bout (Methods). For roughly half of classifiers, errors were dominated by annotation jitter (**Fig 2F-G**). Because they apply a fixed classification rule, HiDRA classifiers may in these cases provide a more reproducible threshold for behavior onset than human observers.

To identify experimental factors that best predict classifier performance, we next compiled a set of candidate features that could impact classifier accuracy, regressed the 82 classifier F1 scores against them, and ranked features by permutation importance (**Fig 2H**). Training data volume was the strongest driver of model performance. An additional strong predictor was behavior rarity, defined as the fraction of scored frames that were labeled for the subject behavior, suggesting that data imbalance corrections(*53*) could improve performance. Finally, behaviors with shorter bouts also had lower F1 scores, likely reflecting annotation jitter at bout boundaries.

Notably, the video framerate, video resolution, and number of tracked keypoints were only weakly correlated with classifier performance, and were not predictive of performance in the permutation importance test. This suggests that within the broad range of experimental settings represented in DooM, HiDRA classifier performance is largely invariant to acquisition hardware and tracking configurations.

### HiDRA classifiers generalize across experimental settings

HiDRA’s high performance in diverse experimental conditions shows that its learned representation is highly expressive, but does not tell us how it is structured. If HiDRA entangled movement with lab-specific appearance, classifier heads for the same behavior would need distinct readout weights across labs, whereas if it learned a shared representation of movement, those heads would be similar. Supporting the latter, cosine similarity among HiDRA heads is high for the same behavior across labs (**Suppl Fig 4**).

Because HiDRA is trained on data from many labs, and appears to have learned a common representation of movement, we hypothesized that HiDRA classifiers could also be transferred between labs without retraining. To test this, we first evaluated HiDRA on the CalMS21 benchmark. Although CalMS21 pose trajectories were included in training HiDRA’s self-supervised GNN, CalMS21 behavior annotations were never seen by the supervised classification network. Strikingly, all tested HiDRA classifiers agreed closely with held-out human annotations on CalMS21 (**Fig 3A**), and achieved competitive F1 scores, despite having been trained on labels from different annotators, on different videos, from different labs (**Fig 3B**).

**Figure 3.**
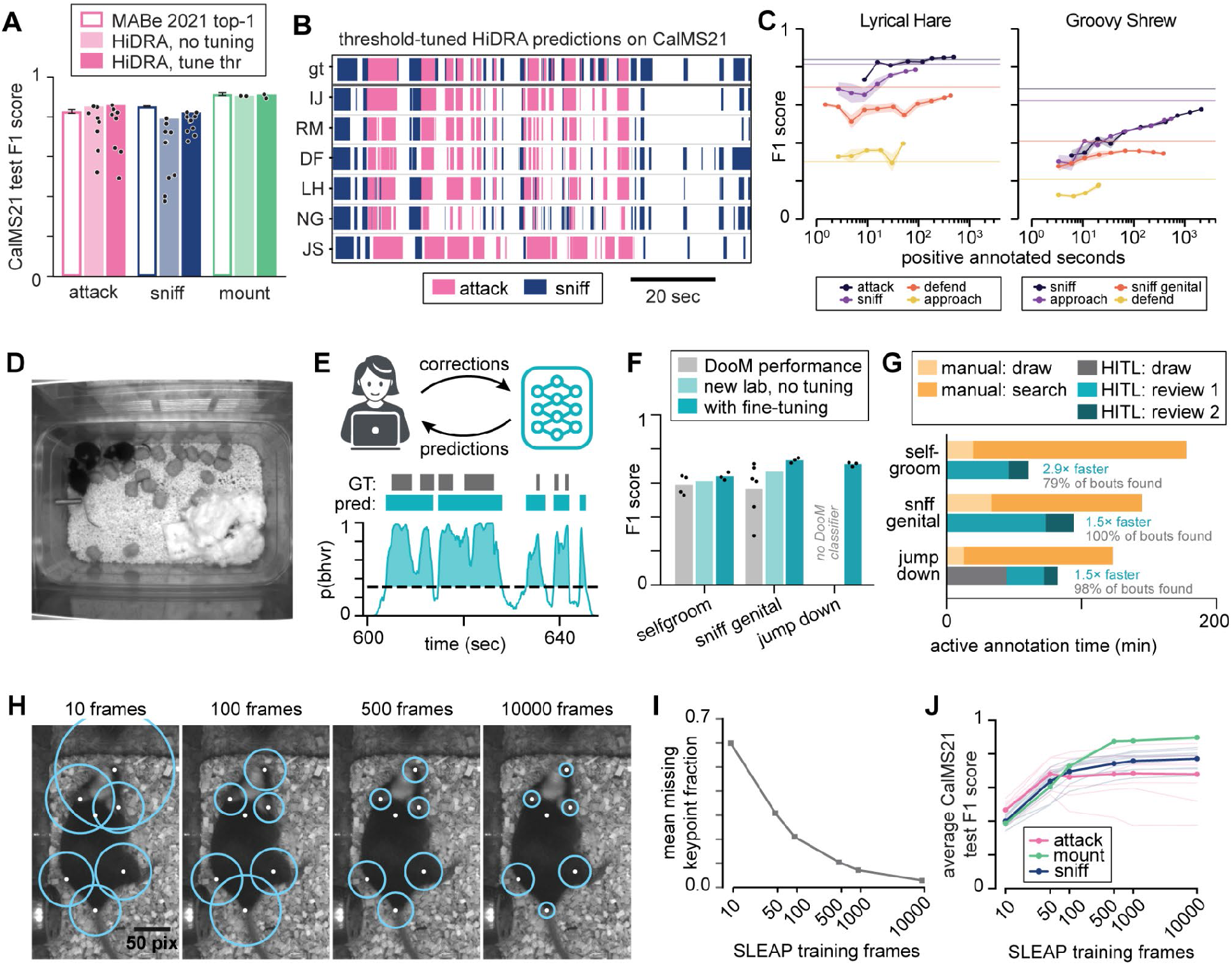
Cross-lab generalization and comparison. **A)** F1 score for attack, investigation (“sniff”), and mounting by HiDRA classifiers for all labs that scored those behaviors (markers) and best-performing classifier (bar), both naive and with threshold fine-tuning, compared to F1 scores achieved by the top-performing model in the original CalMS21 benchmark. **B)** Sample ethogram showing ground-truth manual annotations from CalMS21 (“gt”) alongside predictions from HiDRA classifiers trained on six DooM labs, with threshold tuning on CalMS21 training set. **C)** Fine-tuning of HiDRA classifiers on held-out labs in the leave-one-lab-out experiment, showing F1 score as a function of training set size; dashed lines indicate performance of baseline HiDRA model. **D)** Sample video frame from new lab test dataset. **E)** Schematic of human-in-the-loop (HITL) classifier training (above) and threshold tuning (below). **F)** Performance of HiDRA classifiers on DooM labs (reproduced from Figure 2) compared to performance on the new lab’s data, out-of-the-box (“no tuning”) and after fine-tuning. Plotted points for DooM bars reflect F1 score of different labs’ classifiers on their test sets; plotted points for fine-tuning reflect multiple seeds. **G)** Effort allocation during manual vs HITL behavior annotation. **H)** 90% accuracy radii of SLEAP models trained on the CalMS21 benchmark, with increasing amounts of training data. **I)** Fraction of missing keypoints in SLEAP predictions as a function of SLEAP training set size. **J)** F1 score of HiDRA classifiers on pose-degraded CalMS21, threshold tuning only (thick lines = averages across classifiers: 8 labs for attack, 10 labs for sniff, 2 labs for mount+intromit, collectively labeled here as mount.)

To test broader generalizability, we performed a leave-one-lab-out experiment on DooM, retraining the HiDRA supervised classification network with all data from one held-out lab excluded. F1 scores on held-out labs were variable: in home-cage settings with typical pose skeletons, performance neared that of the base model. In settings that were unique to single labs, such as four-animal experiments and datasets with unconventional pose skeletons, F1 scores were lower, although they could be recovered with fine-tuning (**Fig 3C, Suppl Fig 5**).

Finally, we examined cross-lab performance within DooM itself, by evaluating each HiDRA classifier on every lab that annotated that behavior. In keeping with the leave-one-lab-out lab experiment, some classifiers performed as well on other labs’ data as on their own (**Suppl Fig 6-7**). However, other classifiers failed to generalize: this could reflect either technical barriers, like limited training data on some tasks or a failure of those classifiers to learn lab-invariant readouts, or more biological ones, like lab-to-lab differences in how behaviors are operationally defined or how animals perform the behavior.

### HiDRA classifiers reduce annotation burden for both pose and behavior

To support training of new HiDRA classification heads, we developed and benchmarked a human-in-the-loop (HITL) classifier training pipeline using a new dataset of three-animal homecage behavior that was never used in HiDRA’s development (**Fig 3D**). In HITL annotation, a small number of manual behavior annotations are used to train an initial machine learning model; predictions of that model on held-out data are then manually refined, and training is iterated (**Fig 3E**).

We annotated three behaviors, two of which (self-groom and sniff genital) had pretrained HiDRA classifiers. In line with our generalization experiments, HiDRA’s self-groom and sniff genital classifiers performed well before any fine-tuning (**Fig 3F**). We then fine-tuned only the output head of each classifier on annotations from two other clips. This took under 10 minutes on a single GPU and improved both classifiers (Methods). The same procedure produced an accurate classifier for a new behavior (jump down) by retraining the head of the closest pretrained class, climb, which detected jump down at near-chance level (F1 = 0.07).

Importantly, head fine-tuning leaves HiDRA’s other classifiers untouched, so a new behavior can be added in minutes without affecting the rest of the model. Training additional layers of the supervised model (temporal filtering layers plus classifier head, **Fig 2A**) took over an hour and did not improve performance (**Suppl Fig 8**). Manually annotating these three behaviors in each of three mice in 30 min of video took 8.7 hours, with a majority of that time spent searching the video for bouts. HITL replaces this search with a review of HiDRA’s candidate bouts, which cut annotator time 1.9- to 4.5-fold (**Fig 3G**, Methods).

The second annotation bottleneck faced by HiDRA users is in training pose models. Therefore, we next examined the dependence of classifier performance on pose model training set size. Using the manual pose annotations from CalMS21, we trained SLEAP pose models on between 10 and 10,000 frames (*32*). Smaller training sets produced more dropped keypoints and greater pose jitter, producing a range of structured pose errors (**Fig 3H**). Whereas pose estimation quality continued to improve at up to 10,000 frames of training data (**Fig 3I**), zero-shot HiDRA performance nearly plateaued by 100 frames for attack and sniff and 500 frames for mounting (**Fig 3J**), indicating that HiDRA is highly robust to pose estimation errors. This demonstrates its compatibility with practical usage in new experimental settings with minimal annotation burden.

### HiDRA enables interpretable high-throughput behavioral phenotyping

To demonstrate how HiDRA can be used for behavioral study, we tested its classifiers on a previously published dataset(*24*) of resident-intruder behavior in males from four mouse lines with autism-associated mutations (**Fig 4A**). HiDRA classifiers reproduced previously reported phenotypes of these animals with no fine-tuning (**Fig 4B**, **Suppl Fig 9**), and uncovered further differences in other behavior categories (**Fig 4C-D**).

**Figure 4.**
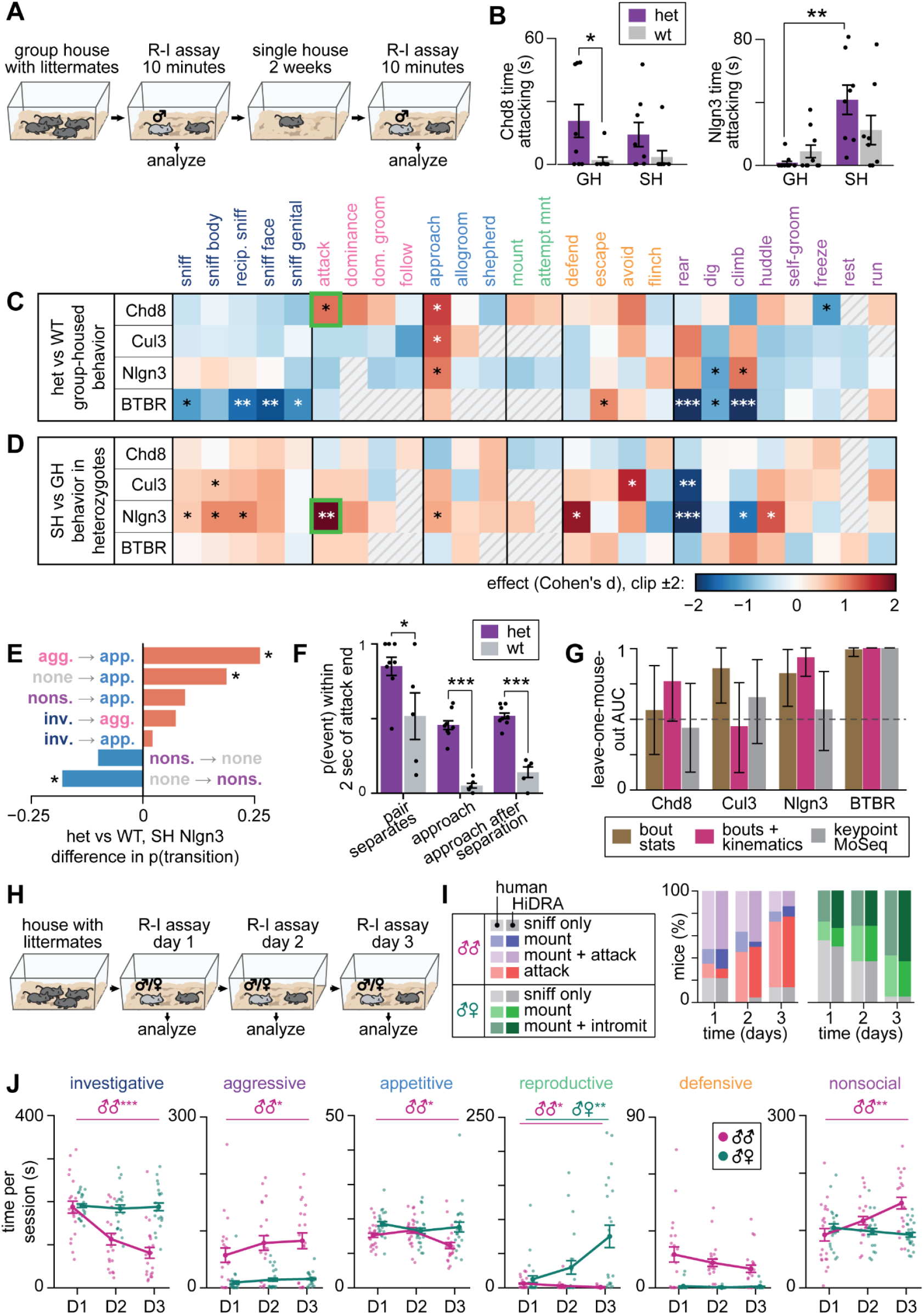
Application of HiDRA to sample experimental datasets. **A)** Design of the autism-line experiment Segalin et al. (1). **B)** HiDRA-derived time spent attacking in group-housed animals, showing that HiDRA classifiers recapitulate two statistical significance findings of Seglin et al. (n=8-10 mice per condition, exact group sizes in Methods; mean ± SEM) **C)** Effect of genotype on time spent in each of the 25 behaviors in group-housed mice, expressed as Cohen’s d. Behaviors are grouped and colored by category. Stars indicate significance from two-tailed t-tests. Hatched cells could not be tested because no animal expressed the behavior in either the heterozygous or wild-type group. Green box marks the comparison shown in B. **D)** Effect of housing in het mice (SH vs. GH within mice), expressed as Cohen’s d. Stars indicate significance from paired t-tests. Green box marks the comparison shown in B. **E)** Behavior transitions that show elevated (red) or reduced (blue) likelihood in heterozygous Nlgn3 mice compared to wild-type littermates, both groups tested in the single-housed condition. F) Enhanced likelihood of attack-approach transitions in heterozygotes can be explained in terms of an increased likelihood of separation following attack in heterozygotes, as well as an increased likelihood of re-engagement within a two-second window. (n=8 mice per condition, mean ± SEM). **G)** Performance of decoder predicting genotype of a held-out mouse using behavior bout statistics produced by HiDRA, statistics + kinematics, or pure unsupervised movement segmentation (see Methods). **H)** Design of the multiday experience experiment. Group-housed male mice were tested in an R-I assay over three days with either a male or female intruder (n=22 paired with male, n=19 paired with female). **I)** HiDRA-derived statistics compared to manual labeling: percent of mice showing mounting and attack behaviors towards male intruders (left) and mounting and intromission behaviors towards female intruders (right), as a function of experience in the resident-intruder assay. **J)** Time per session spent in each of the six behavior categories on days 1-3 for male-male and male-female pairs. Points are individual mice; lines show mean +/-SEM. Stars mark paired t-test on the per mouse day 3 - day 1 difference within each pairing type, Benjamini-Hochberg corrected over seven (object-oriented not shown) behavioral categories (* q < 0.05, ** q < 0.01, *** q < 0.001).

HiDRA can also be used to identify changes in the transition structure of behaviors. For example, computing the transition matrix for single-housed Nlgn3 heterozygotes and their wild-type littermates revealed several transition types that showed enrichment or reduction in heterozygotes (**Fig 4E**). Noting the enhanced probability of transition from aggressive to appetitive behaviors, we examined behavior following the termination of a fighting bout, revealing that the increased rate of attack to approach transitions stemmed from heterozygotes showing an increased propensity to separate from and re-engage social partners (**Fig 4F**). Thus, beyond scoring how much time animals spend engaging in behaviors, HiDRA annotations can be combined with interpretable pose-derived features to gain an understanding of detected phenotypes.

To evaluate the information content of HiDRA annotations and pose data, we trained decoders to infer the genotype of mice in the autism dataset (**Fig 4G**). Bout statistics alone were sufficient to distinguish genotypes in two of four lines. Incorporating behavior-conditioned movement features outperformed bout statistics on every line, achieving performance that was comparable to decoders trained on keypoint-MoSeq syllables(*54*). HiDRA-conditioned movement features are therefore a phenotype readout on par with a purpose-fitted unsupervised model, but are obtained without model training and are natively human-interpretable.

In a second dataset, we tested how behavior changes with repeated testing alone: 41 male mice tested on three consecutive days with either male or female intruders(*55*) (**Fig 4H**). These recordings were part of the self-supervised training of HiDRA, but not of the classification heads. We found that HiDRA classifiers closely agreed with manual annotators in scoring effects of experience, including those driven by subtle distinctions such as mounting with vs without intromission (**Fig 4I**). Extending our analysis to the full suite of HiDRA classifiers, we found a broad, intruder-sex-specific restructuring of resident behavior across days, spanning appetitive, investigative, aggressive, and nonsocial behaviors (**Fig 4J**).

## Discussion

Automated behavior phenotyping has long promised to remove bottlenecks in behavioral neuroscience, scaling assays both in resolution and throughput. Progress has been constrained by a structural problem: building a classifier that generalizes across experimental contexts and research sites requires expertise in both animal behavior and machine learning. The result has been a field of one-off tools, each trained and deployed within individual labs. By assembling fifteen laboratories into a consortium willing to share raw, previously siloed data, we established a dataset substantially larger and more representative of real world variability than was previously available (**Suppl. Table 1**). This data curation effort opened automated phenotyping as a challenge for the machine learning community, and a Kaggle-hosted competition drew 1,781 participants from 85 countries to explore this problem in a span of two months. We release DooM’s videos, pose tracking, and behavior annotations openly alongside HiDRA to facilitate further development.

DooM’s scale and heterogeneity enabled us to ask a question that could not be studied previously, namely whether classifiers trained across many laboratories can learn generalized representations of animal movement. It is possible to treat each DooM lab’s data as a separate classification task, and even a single model trained on the full DooM dataset might have solved the classification problem by learning lab-specific solutions. But we instead found that HiDRA encodes the same behavior from different labs in a similar way (**Suppl Fig. 4**), to the extent that the model can be transferred to laboratories and datasets it was never trained on (**Fig 3**). This distinguishes HiDRA from previous supervised behavior classifiers, which by construction cannot be shared across labs, and for the first time provides a platform for true behavioral generalizability and reproducibility across research sites.

Perhaps the most unexpected finding to emerge from DooM is what did not predict classifier performance. We had expected data acquisition parameters like frame rate, resolution, and number of tracked keypoints to be major drivers of performance. Instead, these factors were only weakly associated with F1 score, and were not predictive in a permutation importance analysis (**Fig 2H**). This argues against the instinct to over-invest in acquisition hardware: a lab does not need a high-framerate, high-resolution camera, a densely tracked skeleton, or a video foundation model (**Suppl Fig 3**) to achieve accurate classification of rodent behaviors. Thus, democratization of behavioral classification is accessible to all labs regardless of hardware and compute constraints.

To demonstrate how HiDRA can support biological insight, we used HiDRA to reanalyze resident-intruder data from four autism-associated mouse lines and from a multi-day experience assay, applying all 33 behavior classifiers simultaneously. Our analysis revealed genotype-, housing-, and experience-specific effects (**Fig 4**). Because each HiDRA head is a named behavior rather than a latent dimension, the effects in **Fig 4** are human-interpretable, simplifying the transition from data visualization to design of follow-up experiments. HiDRA’s utility should extend to any setting where internal state or experimental context might reshape social behavior, with applications in drug screening, circuit perturbation experiments, addiction, metabolic research, and aging studies.

### Limitations

While HiDRA characterizes whole-body social behavior, it is not a substitute for classifiers targeting fine motor phenotypes such as gait or grasping. It cannot report behavior outside its 82 named heads unless a new classifier is trained. An open-vocabulary approach such as MouseGPT(*41*) could in principle identify additional behaviors, but will require careful vetting. Additionally, while HiDRA is robust to noisy pose estimates, it cannot correct identity swaps. We note however that unexpected HiDRA labels could be useful as an indicator that swaps have occurred: for example, an unusually low F1 score on one DooM video helped us find and correct an identity swap in that video’s tracking. Increasing the output of behavior assays from one value to 82 also increases the importance of appropriate correction for multiple comparisons. Users of HiDRA will need to select a correction appropriate to their experimental design. HiDRA does not yet report per-bout confidence scores, and it is best used to augment manual annotation rather than to fully replace it. Finally, HiDRA has so far only been validated in mice, and its extension to other small quadrupeds remains to be tested.

Creation of HiDRA was enabled by DooM, which offers many opportunities for further study and extension. However, its scale and heterogeneity, and the heterogeneity of behavioral experiments in general, still present barriers to growth. Harmonization of the 15 contributed datasets included data intake, pose label standardization, and curating 121 raw labels to 33 named behaviors, as well as collecting experimental metadata for which no consensus standards exist. The list of post-competition corrections (Methods) speaks to the many subtle ways that data intake can fail. Extending this dataset while maintaining quality is nontrivial, and will require sustained effort by domain experts.

## Data and Code Availability

General information about HiDRA may be found at: https://neuroethology.github.io/hidra-site

The DooM dataset may be found on the EMBER archive at: dandi.emberarchive.org/dandiset/000564?pos=1

The competition version of the dataset, which excludes videos and the corrections outlined in the Methods, may be found at: http://kaggle.com/competitions/MABe-mouse-behavior-detection/.

Code for running HiDRA may be found at: https://github.com/neuroethology/HiDRA

Code for human-in-the-loop annotation and classifier training may be found at: https://github.com/talmolab/laras-labeler

The weights for the trained HiDRA model may be found at: https://huggingface.co/Neuroethology/HiDRA

## Data contributors

David J. Anderson¹, Eiman Azim², Dorian Battivelli³^,†^, Daniel W. Bayless⁴, Camilla Bellone⁵, Asaf Benjamin⁶, Joeri Bordes⁷, Ioana Carcea⁸, Alon Chen⁶, Michael J. Conoscenti⁹, Bing Dai¹⁰, Elise Davis¹¹, Kaela Evans¹¹, Annegret L. Falkner¹², Benoit Girard⁵, Nastacia L. Goodwin¹³, Ryan Gorzek¹⁴, Cornelius T. Gross³, Chongye Han¹¹, Weizhe Hong¹⁵, Marcus Intal¹¹, Jorge M. Iravedra-García¹², Kentaro K. Ishii¹⁶, Tomomi Karigo¹⁷, Dayu Lin¹⁸, Sebastián Malagón Pérez¹⁹, Mariela Mendoza¹¹, Simon R.O. Nilsson²⁰, Nguyen Phi¹⁴, Tara Raam¹⁴, Carlos Robles¹¹, Mathias V. Schmidt⁷, Nirao M. Shah²¹, Yair Shemesh⁶, Steven A. Siegelbaum²², Stefanos Stagkourakis²³, Garret D. Stuber¹⁶, Andres Villegas²¹, Lindsay Willmore¹², Ofer Yizhar⁶, Moriel Zelikowsky²⁴, Xingjian Zhang¹⁴

## Kaggle top solutions

Ahmet Erdem²⁵, Daniel Krstevski²⁶, Yudai Ogura²⁷, Matheus Pedroza²⁶, Eduardo Rocha de Andrade²⁵, Takato Yoshikawa²⁸

## Affiliations

1. Division of Biology and Biological Engineering, Tianqiao and Chrissy Chen Institute for Neuroscience, and Howard Hughes Medical Institute, California Institute of Technology, Pasadena, CA, USA
2. Molecular Neurobiology Laboratory, Salk Institute for Biological Studies, La Jolla, CA, USA; Department of Neurobiology, School of Biological Sciences, University of California San Diego, La Jolla, CA, USA
3. Epigenetics and Neurobiology Unit, European Molecular Biology Laboratory (EMBL) Rome, Monterotondo, Italy
4. Salk Institute for Biological Studies, La Jolla, CA, USA
5. Department of Basic Neurosciences, Faculty of Medicine, University of Geneva, Geneva, Switzerland
6. Department of Brain Sciences and Department of Molecular Neuroscience, Weizmann Institute of Science, Rehovot, Israel
7. Research Group Neurobiology of Stress Resilience, Max Planck Institute of Psychiatry, Munich, Germany
8. Department of Pharmacology, Physiology and Neuroscience, Rutgers New Jersey Medical School, Newark, NJ, USA; Brain Health Institute, Rutgers University, Piscataway, NJ, USA
9. Department of Neurobiology, Spencer Fox Eccles School of Medicine, University of Utah, Salt Lake City, UT, USA
10. Neuroscience Institute, Department of Neuroscience and Physiology, New York University Grossman School of Medicine, New York, NY, USA
11. Computational Neurobiology Lab, Salk Institute for Biological Studies, La Jolla, CA, USA
12. Princeton Neuroscience Institute, Princeton University, Princeton, NJ, USA
13. Department of Neurobiology & Biophysics, University of Washington, Seattle, WA, USA
14. Department of Biological Chemistry and Department of Neurobiology, David Geffen School of Medicine, University of California, Los Angeles, Los Angeles, CA, USA
15. Department of Biological Chemistry, Department of Neurobiology, and Department of Bioengineering, University of California, Los Angeles, Los Angeles, CA, USA
16. Center for the Neurobiology of Addiction, Pain, and Emotion; Department of Anesthesiology and Pain Medicine; and Department of Pharmacology, University of Washington, Seattle, WA, USA
17. Kennedy Krieger Institute, Solomon H. Snyder Department of Neuroscience, Johns Hopkins University School of Medicine, Baltimore, MD, USA
18. Institute of Translational Neuroscience, Department of Neuroscience, and Department of Psychiatry, New York University Grossman School of Medicine, New York, NY, USA; Center for Neural Science, New York University, New York, NY, USA
19. Department of Neuroscience, Northwestern University, Chicago, IL, USA
20. Netholabs, Division of Data Science, London, England
21. Department of Psychiatry and Behavioral Sciences and Department of Neurobiology, Stanford University School of Medicine, Stanford, CA, USA
22. Departments of Neuroscience and Pharmacology, Zuckerman Mind Brain Behavior Institute, Vagelos College of Physicians and Surgeons, Columbia University, New York, NY, USA
23. SciLifeLab and Department of Neuroscience, Karolinska Institutet, Stockholm, Sweden
24. Department of Neurobiology and Howard Hughes Medical Institute, University of Utah, Salt Lake City, UT, USA
25. NVIDIA
26. Independent researcher
27. ZOZO Inc., Chiba, Japan
28. DeNA Co., Ltd., Tokyo, Japan

†Current address: Life Sciences Centre (LSC), Vilnius, Lithuania

## Acknowledgements

We thank Addison Howard, Sohier Dane, and Elizabeth Park at Kaggle for their support in coordinating the Days of our Mice Kaggle Competition, and members of the Kennedy lab for comments and feedback during project development.

This work was supported by the Kaggle Research Grant Program, and by NIDA R01 DA059374 and P30 DA048736 (to S.G.), NSF DMR-2433348, NSF IIS-2505098, NVIDIA Academic Grant Program, and Bowers Undergraduate Research Experience at Cornell program (to J.J.S.), NIMH RF1 MH132653, NIDA U01 DA063581, Project ALS, and Salk Collaboration Grant (to T.P.), and NIH Brain Initiative RF1 NS132912, U01 NS131406, Pew Biomedical Scholars Award, and McKnight Scholar Award (to A.K.).

Work of laboratories in the Days of our Mice Consortium is supported by NIH R01 MH070053 (to D.J.A.); EMBL EIPOD4 fellowship (to D.B.); Swiss National Science Foundation 310030_212219 and ERC Consolidator Grant 864552 (to C.B.); NIH K99/R00 106744 (to I.C.);Vera and John Schwartz Professorial Chair in Neurobiology, Ruhman Family Laboratory for Research in the Neurobiology of Stress, the Licht Family (to A.C.); NIH F31 MH125587, NIH T32 NS099578, Washington Research Foundation Postdoctoral Fellow (to N.L.G.); EMBL core funding, ERC Advanced Grant TERRITORY 101097411 (to C.T.G); NIH R01 NS113124, R01 MH130941 (to W.H.); NIH R01 MH101377 R01 MH124927 (to D.L.); NIH F32 MH123049, K99 MH133159, BBRF Young Investigator Grant 30409 (to T.R.); German Research Foundation (DFG) SCHM 2360/12-1, SCHM 2360/8-1, ERA-Net EU 01EW2501B (to M.V.S.); NIH R01 NS049488, R01 NS083872 R01 HD108363 (to N.M.S.); NIH R01 MH120292 (to S.A.S.); NIH R01 DA054317, R01 DA038168, R37 DA032750 (to G.D.S.) and NIH P30 DA048736 (to University of Washington); ERC Advanced Grant NEST 101267597, Joseph and Wolf Lebovic Charitable Foundation Chair for Research in Neuroscience (to O.Y.); NIH R00 MH108734, R01 MH132822, R01 HD117924, and McKnight Scholar Award (to M.Z.)

## List of Supplementary Materials

Days of our Mice Consortium members and affiliations

## Materials and Methods

Figs S1 to S9

