## Supplementary material for "A community benchmark enables high-resolution, cross-lab quantification of mouse social behavior": Methods and Supplementary Material

**List of Supplementary Materials**

Days of our Mice Consortium members and affiliations

Materials and Methods

Figs S1 to S9

### The Days of our Mice Consortium

#### Data contributors

David J. Anderson[1], Eiman Azim[2], Dorian Battivelli[3,†], Daniel W. Bayless[4], Camilla Bellone[5], Asaf Benjamin[6], Joeri Bordes[7], Ioana Carcea[8], Alon Chen[6], Michael J. Conoscenti[9], Bing Dai[10], Elise Davis[11], Kaela Evans[11], Annegret L. Falkner[12], Benoit Girard[5], Nastacia L. Goodwin[13], Ryan Gorzek[14], Cornelius T. Gross[3], Chongye Han[11], Weizhe Hong[15], Marcus Intal[11], Jorge M. Iravedra-García[12], Kentaro K. Ishii[16], Dayu Lin[17], Sebastián Malagón Pérez[18], Mariela Mendoza[11], Simon R.O. Nilsson[19], Nguyen Phi[14], Tara Raam[14], Carlos Robles[11], Mathias V. Schmidt[7], Nirao M. Shah[20], Yair Shemesh[6], Steven A. Siegelbaum[21], Stefanos Stagkourakis[22], Garret D. Stuber[16], Andres Villegas[21], Lindsay Willmore[12], Ofer Yizhar[6], Moriel Zelikowsky[23], Xingjian Zhang[14]

#### Kaggle top-five solutions

Ahmet Erdem[24], Daniel Krstevski[25], Yudai Ogura[26], Matheus Pedroza[25], Eduardo Rocha de Andrade[25], Takato Yoshikawa[27]

#### Affiliations

1. Division of Biology and Biological Engineering, Tianqiao and Chrissy Chen Institute for Neuroscience, and Howard Hughes Medical Institute, California Institute of Technology, Pasadena, CA, USA
2. Molecular Neurobiology Laboratory, Salk Institute for Biological Studies, La Jolla, CA, USA; Department of Neurobiology, School of Biological Sciences, University of California San Diego, La Jolla, CA, USA
3. Epigenetics and Neurobiology Unit, European Molecular Biology Laboratory (EMBL) Rome, Monterotondo, Italy
4. Salk Institute for Biological Studies, La Jolla, CA, USA
5. Department of Basic Neurosciences, Faculty of Medicine, University of Geneva, Geneva, Switzerland
6. Department of Brain Sciences and Department of Molecular Neuroscience, Weizmann Institute of Science, Rehovot, Israel
7. Research Group Neurobiology of Stress Resilience, Max Planck Institute of Psychiatry, Munich, Germany
8. Department of Pharmacology, Physiology and Neuroscience, Rutgers New Jersey Medical School, Newark, NJ, USA; Brain Health Institute, Rutgers University, Piscataway, NJ, USA
9. Department of Neurobiology, Spencer Fox Eccles School of Medicine, University of Utah, Salt Lake City, UT, USA
10. Neuroscience Institute, Department of Neuroscience and Physiology, New York University Grossman School of Medicine, New York, NY, USA

11. Computational Neurobiology Lab, Salk Institute for Biological Studies, La Jolla, CA, USA
12. Princeton Neuroscience Institute, Princeton University, Princeton, NJ, USA
13. Department of Neurobiology & Biophysics, University of Washington, Seattle, WA, USA
14. Department of Biological Chemistry and Department of Neurobiology, David Geffen School of Medicine, University of California, Los Angeles, Los Angeles, CA, USA
15. Department of Biological Chemistry, Department of Neurobiology, and Department of Bioengineering, University of California, Los Angeles, Los Angeles, CA, USA
16. Center for the Neurobiology of Addiction, Pain, and Emotion; Department of Anesthesiology and Pain Medicine; and Department of Pharmacology, University of Washington, Seattle, WA, USA
17. Institute of Translational Neuroscience, Department of Neuroscience, and Department of Psychiatry, New York University Grossman School of Medicine, New York, NY, USA; Center for Neural Science, New York University, New York, NY, USA
18. Department of Neuroscience, Northwestern University, Chicago, IL, USA
19. Netholabs, Division of Data Science, London, England
20. Department of Psychiatry and Behavioral Sciences and Department of Neurobiology, Stanford University School of Medicine, Stanford, CA, USA
21. Departments of Neuroscience and Pharmacology, Zuckerman Mind Brain Behavior Institute, Vagelos College of Physicians and Surgeons, Columbia University, New York, NY, USA
22. SciLifeLab and Department of Neuroscience, Karolinska Institutet, Stockholm, Sweden
23. Department of Neurobiology and Howard Hughes Medical Institute, University of Utah, Salt Lake City, UT, USA
24. NVIDIA
25. Independent researcher
26. ZOZO Inc., Chiba, Japan
27. DeNA Co., Ltd., Tokyo, Japan

† Current address: Life Sciences Centre (LSC), Vilnius, Lithuania

### Materials and Methods

### Contributed datasets

#### Dataset collection.

We cleaned and assembled 15 video and manual behavior annotation datasets from 14 contributing labs. Datasets were required to include roughly six or more hours of top-view video of interacting mice, with manual, frame-by-frame annotations for one or more behaviors. For each dataset, contributing labs completed a datasheet of experimental metadata, including information about the animal, the arena, how videos were acquired, how pose estimation was performed, and any criteria for data exclusion. The datasheet format was adapted from the Datasheets For Datasets standard(*1*), with custom behavioral and experimental metadata questions designed by us based on metadata published for our previous benchmark datasets(*2*, *3*). While some datasets have been part of previously published papers, none of the raw video, behavior annotation, or pose estimation files have previously been released. We converted all pose estimates and behavior annotations to a common format.

Contributed videos were all derived from previously conducted experiments and reflected a range of standard and specialized assays of animal social behavior. **Table 1** summarizes the source experiments for each lab; note that datasets from some labs were compiled across multiple experimental settings. Due to the large file size of the video dataset and the cost of training video-based models, challenge participants were only given access to behavior annotations, pose estimates, and experimental metadata, not raw video files.

| Dataset | Contents | Camera | Pose tracking | Behavior annotation |
|---|---|---|---|---|
| AdaptableSnail | Longitudinal group monitoring in enriched environment | Sony SNC-VB770 4k network camera | DeepLabCut v2.2 / 2.3 (7 models) | Media Player Classic v1.9.12 + Microsoft Excel 2016 |
| BoisterousParrot | Virgin/dam shepherding assay | Blackrock Microsystems IR camera | DeepLabCut v2.0 | BORIS |
| CautiousGiraffe | Free social interactions, same- and opposite-sex pairs | Logitech C920 | DeepLabCut multi-animal v2.3 | Noldus EthoVision XT16 |
| DeliriousFly | Male resident/intruder assay | Imaging Source camera with wide angle TPL 0420 6MP lens | DeepLabCut v2.2 | BORIS |
| ElegantMink | Female mating behavior assay<br>Male mating behavior assay<br>Male resident/intruder assay | ELP 720p USB camera | SLEAP v1.3.3 | Custom MATLAB program |
| GroovyShrew | Female homecage social behavior | ELP 1.3 MP USB camera | SLEAP v1.2.1 | Caltech Behavior Annotator |
| InvincibleJellyfish | Experience-dependent resident/intruder assay | Basler acA1300-75gm | MARS | Noldus Observer |
| JovialSwallow | Male resident/intruder assay | unknown | SLEAP v1.3.3 | Noldus EthoVision XT12 |

| | | | | |
|---|---|---|---|---|
| LyricalHare | Social defeat stress assay | FLIR | SLEAP v1.2.2 | BORIS |
| NiftyGoldfinch | Neutral arena social investigation assay | Webcam | SLEAP v1.1.5 | Caltech Behavior Annotator |
| PleasantMeerkat | Two-chamber territorial behavior assay | Basler acA2040-55uc | DeepLabCut multi-animal v2.2 | Solomon Coder beta.19.08.02 |
| ReflectiveManatee | Male resident/intruder assay | Basler acA640-300gc | DeepLabCut v2.3.0 | Caltech Behavior Annotator |
| SparklingTapir | Operant conditioning aggression assay<br>Chronic social defeat stress assay<br>Male resident/intruder aggression assay | Basler acA2040-120uc; WhiteMatter e3Vision | DeepLabCut (3 models) | SimBA annotation interface |
| TranquilPanther | Female resident/intruder assay<br>Female mating behavior assay | Basler | DeepLabCut v2.2<br>SLEAP v1.3.3 | BORIS v7.9.1 |
| UppityFerret | Neutral arena social investigation assay | unknown | DeepLabCut v2.2b7 | SimBA annotation interface |

**Table 1. DooM source data.** the social assays, cameras (where known), tracking software, and annotation software used in the collection of the DooM dataset. Further information about each experiment is provided in the **DooM Datasheet**.

#### Animal background.

For full details of animal strains, housing, and experimental conditions, see the **DooM Datasheet**. An abridged summary of this information is provided in **Table 2**. All US-based experiments were carried out in accordance with the NIH guidelines and approved by the Institutional Animal Care and Use Committee at the host institution. International experiments were conducted under guidance of the host country's animal experimentation law. All experiments were performed under veterinary supervision.

Because a goal of the dataset is to study naturalistic, self-initiated behavior, we did not include data from animals undergoing optogenetic, chemogenetic, or pharmacological perturbations. However, a subset of animals had undergone previous experimental manipulations known to impact behavior, including chronic social isolation, mating experience, fighting experience, litter-bearing, foot shock stress, and social defeat stress. In addition, several datasets contained mice with head-mounted neural recording devices or optogenetic fibers; while animals are typically habituated to the weight of these devices prior to experimentation, the presence of the devices and attached cables (when present) has the potential to impact animal behavior.

| **Dataset** | **Animal background** | **Experimental design** | **Care & ethics** |
|---|---|---|---|

| | | | |
|---|---|---|---|
| AdaptableSnail | Male ICR/CD1 mice; groups of 3–4; bred at Weizmann Institute of Science and Max Planck Institute of Psychology | Group-housed 1-2 wk in enriched “Social Box” arena (60×60 cm; feeders, ramps, S-wall, sheltered/unsheltered nests); some groups regrouped at day 9; wireless head-mounted neural loggers on some groups. Continuous 12 h light/dark cycle in arena under naturalistic light | IACUC + IBC at Weizmann; NIH Guide; veterinary supervision |
| BoisterousParrot | Pup-naive C57Bl/6 virgin females (~8 wk), bred at NYU School of Medicine; co-housed with an experienced dam and her P1 litter | Co-housing of one virgin with dam + litter for 4 consecutive days in a large custom home cage; dam "shepherds" virgin toward nest. Virgins kept isolated from dams and pups until testing; co-housing cage with abundant bedding, food pellets and hydrogel; visible light by day / infrared at night | IACUC at NYU School of Medicine; NIH Guide; veterinary supervision |
| CautiousGiraffe | C57Bl/6J mice, male and female; bred at University of Geneva | Free interaction in open field; both animals pre-habituated to arena, 5 min habituation then 10 min recording under 20–30 lux illumination | conducted under Swiss animal experimentation law; veterinary supervision |
| DeliriousFly | Male C57Bl/6J residents vs BALB/c intruders; bred at Columbia | Resident–intruder in resident home cage, 10 min under red-light illumination; one mouse carries head-mounted neural recording device | IACUC + IBC at Columbia; NIH Guide; veterinary supervision |
| ElegantMink | C57BL/6J and 129/SvEvTac, male and female; bred at Stanford | Male mating: hormonally-primed group-housed B6 female into B6 male home cage, 30 min. Female mating: primed B6 female into 129/SvEvTac male cage. Male aggression: group-housed 129/SvEvTac male into B6 male cage, 15 min. Food and water removed during encounter, infrared illumination | IACUC + IBC at Stanford; NIH Guide; veterinary supervision |
| GroovyShrew | C57Bl/6J; female-female and male-female pairs; bred at UCLA | Novel cage, 5 min habituation, 12–25 min recording under Infrared illumination; both mice have head-mounted neural devices (some tethered, some wire-free); some animals reused across videos | IACUC + IBC at UCLA; NIH Guide; veterinary supervision |
| InvincibleJellyfish | Male C57Bl/6J residents and BALB/c intruders; purchased from Charles River | Resident–intruder in home cage; resident recorded ~2 min alone, then 10 min with intruder. Subset of residents underwent mating experience, a series of inescapable footshocks, or 4 weeks chronic social isolation prior to the assay | IACUC + IBC at Utah; NIH Guide; veterinary supervision |
| JovialSwallow | Male C57Bl/6N; bred at Caltech | Resident–intruder in home cage, 15 min under dim red light, recording started 3–10 s after intruder introduction | IACUC + IBC at Caltech; NIH Guide; veterinary supervision |
| LyricalHare | Male CD1 residents, sexually naive, 12–18 wk, bred at Princeton; male BALB/cJ intruders | Resident–intruder in resident home cage, data subsampled from experiment consisting of 8 days × 3 back-to-back 9-min sessions with different BALB/cJ intruders, each preceded by 1-min | IACUC at Princeton; NIH Guide; veterinary supervision |

| | | | |
|---|---|---|---|
| | purchased from Taconic | baseline during which animals are separated by a plexiglass barrier | |
| NiftyGoldfinch | C57BL/6J, male and female, 8–10 wk, 25–30 g, purchased from Jackson Labs (000664) | Individually housed 3 wk before experiments; 12:12 reversed light cycle tested during dark phase under dark red light; 10-min divided "separation" control session, then 20-min free interaction in a new cage with fresh woodchip bedding. Both animals have head-mounted neural device | IACUC at UCLA; NIH Guide; veterinary supervision |
| PleasantMeerkat | Male C57Bl/6J and CD1; bred at EMBL | Territorial assessment: pairs co-housed 48 h in a large two-compartment arena (each side with central cage with food/water); 30 min scored in two periods (20 + 10 min, 70 min apart), recorded under red light | IACUC at EMBL; NIH Guide; veterinary supervision |
| ReflectiveManatee | Male C57Bl/6J residents, BALB/c intruders; bred at NYU | Resident–intruder in mouse cage, 3 min habituation, 10 min test under infrared illumination; one mouse has head-mounted neural device; most residents in two videos | IACUC + IBC at NYU Langone; NIH Guide; veterinary supervision |
| SparklingTapir | Aggressors (CD1 or CFW, male or female) and subordinates (C57, male or female); bred at UW and Rutgers | Three assays: operant social self-administration, home-cage resident–intruder, chronic social defeat stress; some animals in multiple videos | IACUC + IBC at UW and Rutgers; NIH Guide; veterinary supervision |
| TranquilPanther | Female C57Bl/6J subjects paired with a female B6, male B6, or male Swiss Webster; bred at UW | Female home cage, 5 min habituation, 10–120 min sessions recorded under infrared illumination (sexual behavior with males, social interaction with females); one mouse has head-mounted neural device; same animals across videos | IACUC + IBC at UW; NIH Guide; veterinary supervision |
| UppityFerret | Male C57Bl/6N subjects, male CD1 partners; bred at Max Planck Institute of Psychology | Chronic social defeat stress vs non-stressed subject, 10 min habituation then 10 min with CD1 under 15–20 lux illumination | approved by Government of Upper Bavaria; EU Directive 2010/63/EU; veterinary supervision |

**Table 2. Summary of animal backgrounds and experiments.**

#### Pose estimation.

Eleven contributors had performed markerless pose estimation themselves using SLEAP or DeepLabCut, and provided these pose estimates. To track the remaining four datasets, we trained pose models and performed pose estimation ourselves using either SLEAP or MARS. After standardizing names of body parts, contributed datasets used 17 distinct anatomical keypoints and nine unique keypoint configurations to capture mouse pose (**Supplemental Figure 1**). Only three keypoints were present in all configurations: the two ears and the base of the tail.

#### Behavior annotations.

The list of behaviors annotated in the raw contributed datasets included **121 unique strings**, scored using a variety of annotation software. Contributors provided a written description of criteria followed when annotating for each behavior, which we used alongside visual inspection to standardize annotations to **46 distinct behavior names**; standardized names and the original strings mapped to them are provided in **Table 3**.

After naming standardization, removal of behaviors that were only annotated in small subsets of videos, removal of behaviors whose detection relied on non-tracked body parts (e.g. "tail rattling" when the tail was not tracked) and removal of behaviors with under 1 minute of total positive annotations, we obtained a cleaned list of **33 unique behaviors**.

| Canonical name | Type | Raw labels |
|---|---|---|
| aggressiveinvestigation | Social | agg_investigation |
| **allogroom** | Social | groom, socialgroom, grooming, socialgrooming, allogroom |
| **approach** | Social | approach |
| **attack** | Social | attack, a |
| attackattempt | Social | attack_attempt |
| attemptintromit | Social | struggle to intromit |
| **attemptmount** | Social | attemptmount, mount_attempt, attempt mount |
| **avoid** | Social | avoid |
| **biteobject** | Self-directed | biteobj |
| **chase** | Social | chase, chasing, pursuit_prediction, c, charging, pursuit |
| **chaseattack** | Social | ca |
| circle | Social | circle |
| **climb** | Self-directed | climb, cl |
| contact | Social | social contact |
| **defend** | Social | defend, defensive, fightback, u |
| **dig** | Self-directed | dig, digging |
| disengage | Social | walk_away |
| **dominance** | Social | dominance |
| **dominancegroom** | Social | mounting aggressive grooming, mounting_aggressive_grooming, mounting/ aggressive grooming |
| dominancemount | Social | dom_mount, dom_mount_attempt |
| ejaculate | Social | ejaculate, ejaculation, ejaculation_state, res ejaculate |
| **escape** | Social | run, escape, fl, runaway, run away |
| **exploreobject** | Self-directed | exploreobj |

| | | |
|---|---|---|
| **flinch** | Social | flinch |
| **follow** | Social | f, follow |
| **freeze** | Self-directed | freezing |
| genitalgroom | Self-directed | genital_groom |
| **hunch** | Self-directed | b_huddle |
| **intromit** | Social | intromit, intromission |
| lordosis | Social | lordosis |
| **mount** | Social | mount, mounting, copulation |
| **rear** | Self-directed | rear, rearing, female_rearing, standing against wall, stand, up, r, u |
| **reciprocalsniff** | Social | reciprocal, nose_to_nose |
| reject | Social | rejecting time |
| **rest** | Self-directed | rest |
| **run** | Self-directed | run |
| **selfgroom** | Self-directed | grooming, self-grooming, groom, selfgrooming, self grooming, self_grooming, clean, self-groom |
| **shepherd** | Social | shepherding |
| **sniff** | Social | sniff, interaction, exploration, passive, investigation, int sniff, sniffing, general_sniffing, closeinvestigation, closeinvestigate |
| **sniffbody** | Social | unilateral_body, sniffing body, sniff_body |
| **sniffface** | Social | sniffface, sniffing face, sniff_face |
| **sniffgenital** | Social | anogenital, sniffgenitals, male_anogenital_sniffing, unilateral_anogenital, white mouse anogenital sniffing, white_mouse_anogenital_sniffing, anogenital sniffing, anogenital_sniffing, anogenital_prediction, w_nose_to_tail, b_nose_to_tail, sniff_genital, sniff_genitals |
| **submit** | Social | submissive |
| tailrattle | Social | r |
| thigmotaxis | Self-directed | freezing/thigmotaxis, freezing thigmotaxis, freezing_thigmotaxis, freezing/ thigmotaxis |
| **tussle** | Social | tussling |

**Table 3. Raw list of annotated behaviors in the DooM dataset.** A total of 46 behaviors were annotated, with 121 unique strings. 33 of these behaviors that passed additional quality checks were included in the compiled DooM dataset (indicated in **bold**).

Importantly, despite our standardization, behavioral labels are not always defined consistently across labs. For example, if Lab A annotates all sniffing behavior as “sniff,” while Lab B separates “sniff” from “sniff anogenital,” then the label “sniff” refers to different behavioral definitions in each case. Similar problems of annotation granularity exist for mating behavior--where some labs separate attempted mounting, mounting, and intromission, while other labs classify all three as mounting--and for aggression, which some labs broke down into chasing, grappling, and tussling while others lumped these many actions into a single label of attack. This semantic variation is in addition to inter-annotator variability, which can occur even within a lab and further fuels differences in annotations between labs.

The targets for classification were constructed using these behavior labels plus the identity of the animal performing the behavior and the animal the behavior was directed towards, or “self” for single-animal behaviors such as rearing and self-grooming. Thus, the goal of classification is to predict (agent, target, action) triplets. Behaviors are often annotated asymmetrically between animals in an assay. For example, many datasets were assays of interactions between a subject animal and an “intruder” introduced as a stimulus; often in these assays only the actions of the subject animal were annotated, or different subsets of actions were annotated for subject vs stimulus animal. See “Evaluation” section below for information on how this asymmetry in annotations was handled during classifier assessment. Finally, we note that in the case of rare behaviors, in some videos a behavior was annotated for but never occurred.

---

### Kaggle Competition

#### Data format standardization.

The raw data from DooM contributor labs was produced by a variety of pose estimation and annotation software (Table 1), and came in a diverse array of formats. To make these diverse data types accessible to analysis, we converted all pose and annotation data to a standard format, such that data from all labs were stored in tables with a common set of headers.

Pose data was stored in a five-column table with one entry per keypoint per frame, with values:

- `video_frame`
- `mouse_id`
- `bodypart` - The tracked body part.
- `[x/y]` - The body part's X/Y coordinate position in pixels.

Behavior annotations were stored in a five-column table with one entry per bout, with values:

- `agent_id` - ID of the mouse performing the behavior.
- `target_id` - ID of the mouse targeted by the behavior. Self-directed behaviors have the same agent and target ID.
- `action` - What behavior is occurring.
- `[start/stop]_frame` - The first/last frame of the action (inclusive).

Note that the pose format differs from that used in most tracking software: each row contains data for a single tracked keypoint on one frame of video. This allows data to be loaded in the same way regardless of which body parts were tracked.

In both data types, frames are indexed relative to the start of the corresponding video files. Importantly, in some videos, recording began before one or both animals were introduced, or ended after animals were removed- in these cases, frames without both animals were excluded from pose and annotation files. We saved both data types as parquet files, with one parquet per video. Accompanying the experimental data, we constructed a metadata sheet with one entry per video, with the following fields:

- `lab_id`
- `video_id`
- `mouse[1-4] [strain/color/sex/id/age/condition]`
- `frames per second`
- `video duration (sec)`
- `pix per cm (approx)`
- `video [width/height]`
- `arena [width/height] (cm)`
- `arena shape`
- `arena type`
- `body parts tracked`
- `behaviors labeled` (necessary as some labs did not annotate the same behaviors for all videos.)
- `tracking method`

#### Train/test set split.

We manually split each lab's data into training (70%) and test (30%) sets, with the exception of three labs for which proportions of training and test videos were reversed to test few-shot learning and evaluate classifier data-efficiency: CautiousGiraffe, DeliriousFly, and NiftyGoldfinch. Splits were performed at the level of videos; when animal identity was known, all videos of the same individual mice were assigned together into training or test. In cases where videos from a lab were compiled across multiple distinct recording setups or experimental contexts, such as male-male pairings vs male-female pairings, we preserved the 70/30 train-test split within each context to the extent possible.

For the Kaggle competition, test videos were further split 50/50 at the level of videos to create public and private test sets, a strategy used to deter overfitting. During the competition participants could see their lab-averaged F1 scores on the public test set, and could use this to make model design choices. After the competition closed, the final ranking of submissions was determined via evaluation on the private test set only.

#### Extended training data and additional annotations of CalMS21.

To augment the training data available in the Kaggle competition, we applied the above formatting convention to data from the CalMS21 and MABe22 datasets, and for the subset of CRIM13 videos that had two awake behaving mice.

The CalMS21 dataset includes three classification tasks: standard classification (task 1), annotator style matching (task 2), and few-shot learning (task 3), as well as a large collection of pose-only data (supplemental); we included task 1, task 2, and supplemental data for training in the competition. The original CalMS21 task 1 dataset only included annotations for attack, mount, and sniffing behaviors- however the raw data files used to generate that dataset included other annotated actions, which were merged or filtered to create the competition labels. The supplemental dataset was also annotated for a variety of social behaviors, which were omitted from the original CalMS21 dataset release as their quality was not as thoroughly vetted.

To give access to as many annotation styles and behaviors as possible, we extended the CalMS21 task 1 and supplemental datasets to include these additional annotated behaviors, which included body part-directed sniffing annotations (face/body/anogenital), approach, attempted mounting, intromission, dominance mounting, and genital grooming. In total this modified release of CalMS21 includes 301 videos with new annotations and 101 videos with expanded annotations.

#### Evaluation.

We quantified classifier performance using a nested F1 score, which is preferable to accuracy when evaluating the performance of classifiers on class-imbalanced data such as behavior annotations. For each lab, we first computed for each (agent, target, action) triplet the frame-level True Positive (TP), False Positive (FP), and False Negative (FN) counts over all videos in which that triplet was annotated. We then computed that triplet's F1 score as

$$F1 = 2 * precision*recall/(precision + recall) = 2TP/(2TP + FN + FP)$$

We defined the lab-level F1 score to be the average of all that lab's triplet-level F1 scores; note that all actions are thus equally weighted within a lab-level F1 score, regardless of how frequent they are. Finally, we computed the overall F1 score as the average F1 score across labs, weighting all labs equally.

Because some labs provided annotations for different subsets of behaviors in different videos, classifier performance measures should not penalize models for detecting behaviors that were not annotated for. For example, if a video is only annotated for attack and investigation, classifiers should not be penalized for detecting bouts of grooming that occur but were not scored by the lab. Thus, for each file in the test set, we provided a list of (agent, target, action) triplets to be used during evaluation, and competition teams were not penalized for submitting predictions of actions not included in this list.

#### First-place solution.

A graph neural network over the body parts of two interacting mice, pretrained to predict keypoint motion, feeds a bidirectional LSTM classifier that scores each behavior at every frame. Five such models with different sampling rates and skeleton sizes are ensembled and decoded with per-lab, per-behavior thresholds.

**Feature design.** Features are learned rather than hand-crafted. The feature extractor is a graph neural network (GNN) whose nodes are the tracked body parts of the animal and its interacting partner, evaluated over a window of consecutive frames. Each node is initialized from a learned body-part embedding plus that keypoint's displacements over the preceding one to four time bins, with a learned placeholder for missing keypoints. Each edge between nodes is active only when those keypoints are within a fixed physical radius of each other, and the relative displacement vector is the input to the message function. Four residual blocks follow, each combining three operations: message passing along two edge types (either between body parts of the same animal or between body parts of the two animals); a per-node feed-forward layer; and a unidirectional LSTM that carries each node's state forward in time. This network is trained on all tracking data in a self-supervised manner, to predict each keypoint's next-step displacement.

**Data standardization.** Keypoints are mapped onto a shared body-part vocabulary, coordinates are converted to millimetres, and frame rates are unified by interpolating each window onto a fixed sampling rate. A lab embedding and a per-recording annotation mask handle lab-specific vocabularies. An ensemble of five feature-extractor models is trained, using a range of temporal resolutions and numbers of keypoints, as well as a range of augmentation hyperparameters.

| Ensemble member | Sample rate | Body parts | Max scale augmentation | Max time-dilation augmentation | Variance of keypoint noise augmentation | GNN aggregation radius |
|---|---|---|---|---|---|---|
| 11fps_4bp | 11 | 4 | 1.6 | 1.8 | 3.5 | 125 |
| 15fps_5bp | 15 | 5 | 2.0 | 2.0 | 3.0 | 150 |
| 19fps_6bp | 19 | 6 | 1.5 | 1.7 | 2.5 | 175 |
| 23fps_7bp | 23 | 7 | 1.7 | 1.5 | 2.0 | 100 |
| 27fps_6bp | 27 | 6 | 1.9 | 1.4 | 1.5 | 125 |

**Table 4.** Parameters of the five models in the first-place ensemble.

**Model architecture.** The GNN described above is frozen after self-supervised training, and its representations are passed to a supervised classifier. Node features from all four residual blocks of the GNN are concatenated for agent and partner, passed through a per-body-part feed-forward layer, and summed across body parts to give a single vector per timestep. This is then added to a lab ID embedding that is learned during training. This representation is passed through three residual blocks, each a bidirectional LSTM followed by a feed-forward layer, then passed to a final linear layer that emits an independent logit per timestep for each of 33 behaviors, so co-occurring behaviors are scored separately. A threshold for detection is computed separately for each classifier head. One supervised classifier model is trained for each of the five models in the feature-extractor ensemble.

**Training and inference.** The extractor is pretrained, then the classifier is trained on masked binary cross-entropy with early stopping on a per-lab validation split. Inference averages augmented passes across members, applies per-lab, per-behavior thresholds, and reports the top above-threshold behavior per frame and animal pair.

Additional details of training and inference are outlined in the **HiDRA classifier architecture, training and inference** section below.

#### Second-place solution.

Pose from all labs was mapped to a common 12-keypoint skeleton and converted to pairwise distances and kinematics, then fed to an ensemble of CNN-recurrent and SqueezeFormer models trained as frame-wise multi-label segmenters on pooled multi-lab data.

**Feature design.** For solo actions, input pose features included all pairwise intra-animal bodypart distances, as well as per-body-part velocity and acceleration. For social actions, input features also included cross-animal body-part distances, as well as the solo features of both animals. In addition, the sex, frame-rate scale, lab ID, and candidate action labels were passed as sequence-level features.

**Data standardization.** All tracking data were resampled to 30 fps, using linear interpolation for pose features and nearest-neighbor interpolation for labels. Pose keypoints were reduced to a fixed 12-part canonical skeleton consisting of parts [nose, ear_left, ear_right, neck, body_center, lateral_left, lateral_right, hip_left, hip_right, tail_base, tail_midpoint, tail_tip]. Points not tracked by a lab or absent on a frame were replaced with the mean of all body parts present in that frame.

**Model architecture.** Each model received a fixed-length window of 512 frames, and produced frame-wise multi-label predictions for all candidate actions simultaneously, treating the task as dense, temporal segmentation. Two architecture families were used. The first was a multi-scale temporal convolution front end fed into a sequence model (bidirectional GRU or Transformer) and a linear output layer. The second was a SqueezeFormer encoder applied directly to the input sequence. Ten variants of each model (5 solo, 5 pair), differing in depth and hyperparameters, were trained and ensembled to produce the final prediction.

**Training and inference.** Models were trained with a one-vs-rest binary cross-entropy loss over all actions, with a cosine learning-rate schedule with warmup and an exponential moving average of weights with weight decay. Data augmentation included additive Gaussian noise, time stretching, temporal shifting of the input features (labels held fixed), and left-right body-part flipping. Models were trained with 5-fold stratified cross-validation. Predictions were generated with sliding windows of 512 frames at a stride of 128, with edge frames discarded and overlapping predicted probabilities averaged,  thresholded with lab- and action-specific thresholds, and only the behavior with the highest score was emitted.

#### Third-place solution.

A bidirectional LSTM and XGBoost classifiers, trained on ~60 hand-crafted pose features (distances, headings, speeds, body shape, metadata) from a skeleton harmonized across laboratories, are blended and thresholded per lab and behavior on out-of-fold predictions.

**Feature design.** The recurrent model used 62 per-frame features for each ordered agent–target pair: inter- and intra-animal keypoint distances (including 10-frame lags), ellipse-based body-shape descriptors, each animal's heading and its relation to the other animal, per-animal and relative speeds, the midpoint of the two body centers, and metadata (sex, strain, arena properties, frame rate, laboratory). Continuous features were z-scored or quantile-transformed. The gradient-boosted models used a larger set of pairwise keypoint distances, lagged displacements, rolling position, speed, and curvature statistics, and pairwise interaction measures, pruned per behavior by importance. Features were pruned per behavior by importance, with 10-195 features retrained.

**Data standardization.** Pose data from all laboratories were pooled and coordinates converted from pixels to physical units using each video's arena dimensions, without centering or rotational alignment. Skeletons were harmonized to five keypoints (tail base, both ears, nose (with "head" remapped to nose), and body center (substituted by the hip midpoint or the mean of the other points when untracked)) plus tail tip where available. Missing coordinates were linearly interpolated for the recurrent model and handled natively by the tree models. Frame rate was not resampled: the recurrent model expressed time steps relative to 30 fps and included frame rate as a feature, while the tree models computed speeds in cm/s and rescaled all window and lag lengths by frame rate. Tree models were fitted separately within nine keypoint-configuration sections.

**Model architecture.** The ensemble comprised four recurrent networks and two gradient-boosted tree models. Each network took 256-frame windows (512 in one member) and comprised an 8-dimensional laboratory embedding concatenated to the inputs, a bidirectional LSTM (128 units per direction), dropout (0.3), a 256-unit SELU dense layer with a second laboratory embedding, and a softmax over 33 behaviors plus a no-behavior class, with unannotated behaviors masked out. Tree models were XGBoost one-vs-rest classifiers per section, interaction type, and behavior (learning rate 0.08, depth 8, up to 1000 trees).

**Training and inference.** Networks were trained with Adam and a cosine-restart schedule under 5-fold cross-validation grouped by video, with randomized window offsets and shifted-window averaging at inference. Tree models used early stopping on F1, with Savitzky–Golay-smoothed outputs. Predictions were blended (0.43:0.57), thresholded with per-laboratory, per-behavior thresholds optimized on out-of-fold predictions, and resolved by the highest probability-to-threshold ratio.

#### Fourth-place solution.

For each laboratory, the solution trained both XGBoost classifiers on hand-crafted kinematic features and a dilated convolutional network on standardized multi-animal pose sequences. The best cross-validated model was selected for each lab, with majority voting used for three labs.

**Feature design.** XGBoost models used hand-crafted per-frame features: inter-keypoint distances, velocities, acceleration and jerk, body angle and elongation, angular velocity, grooming-specific features, animal metadata, and distance to arena boundaries, with pairwise models adding cross-animal distances, relative speed and orientation, allogrooming features, and pair metadata. Temporal context was added by concatenating features from neighboring frames (±4 at stride 5) or by windowed statistics (±2 at stride 15). For four labs with heavy missing data, keypoints were linearly interpolated with an indicator flag. The neural network took, per keypoint and 24-frame window, availability statistics, coordinate mean and SD, and distance to the arena edge, with inter-animal distances, speed, heading similarity, and previously labeled behaviors injected after the backbone.

**Data standardization.** Videos were resampled to 30 fps (labels by mode), and keypoints reduced to a common skeleton: nose, ears, body center, and tail base for XGBoost, with missing points derived from hips, lateral keypoints, or head; for the network, lateral→hip and head→nose, dropping headpiece and spine.

**Model architecture.** Per lab, binary XGBoost classifiers were trained per action, separately for single-animal and pairwise behaviors. The neural network was a residual dilated 1-D convolutional network with GroupNorm, ELU, and InstanceNorm3d input, a softmax across animal pairs in some layers, pooled global context, and a gate selecting between direct and max-over-pairs classification.

**Training and inference.** XGBoost used 4-fold grouped cross-validation over three seeds, F1-optimized per-action thresholds, and argmax over threshold residuals. The network used masked, prevalence-weighted binary cross-entropy, trained jointly and then fine-tuned per lab, with rare-action logits scaled by 1.5. Per lab, the best cross-validated model was selected, or majority voting for three labs.

#### Fifth-place solution.

Tracking from all labs was mapped onto a shared six-keypoint skeleton and converted to invariant kinematic features, which a graph neural network, per-animal Squeezeformer encoder, and pairwise classification head used to predict agent–target behaviors per frame. A 23-model ensemble was trained jointly across labs with action-biased sampling and class-weighted focal loss.

**Feature design.** All input features were translation- and rotation-invariant kinematic features. Per-mouse features included velocity, acceleration, jerk, angular velocity, heading, and body-configuration measures (curvature, length, elongation, ear spread, head and tail angles). Inter-mouse features were computed from each animal's perspective toward every other animal: centroid distance and its rate of change, body-part distances (e.g., nose to tail base), relative speed, and approach angle. This gave 58 features per mouse (232 per frame for four animals).

**Data standardization.** A six-point "master skeleton" (nose, left ear, right ear, head center, body center, tail base) was defined from the body parts most commonly tracked across labs, with a fallback hierarchy substituting nearby parts when a keypoint was absent. Coordinates were converted to centimeters using each video's calibration; no adjustments for video framerate were made.

**Model architecture.** A TransformerConv Graph Neural Network was trained jointly on all labs. The model treated animals as nodes, enriching each animal's features with information from cage-mates. Each animal's trajectory was then encoded independently in time by four Squeezeformer blocks (51-frame convolution kernel), with cross-animal GNN layers interleaved in six models. Encoded representations were concatenated into all ordered agent–target pairs and classified per frame by an MLP head with a softmax over behaviors plus "no action"; a Relation Network variant added global cage context. Logits were masked to annotated behaviors per video.

**Training and inference.** Training windows (512 frames) were sampled preferentially from annotated frames, and focal loss ($\gamma = 2.0$) with duration-based class weights addressed imbalance. Models used AdamW with a one-cycle schedule, dropout 0.1–0.35, and FP16. Inference used overlapping windows (stride 256) with averaged probabilities, argmax per frame, segment merging, and a minimum-duration filter. The final ensemble comprised 23 models from several architectural variants.

### HiDRA model creation

#### Post-competition cleanup of the DooM dataset.

Evaluation of competition F1 scores on individual videos and (lab, behavior) pairs revealed several data quality issues missed during the initial dataset screening. We used these to make several corrections to the DooM dataset, which are applied in the official version of the dataset shared publicly. For reproducibility purposes, we have not applied these corrections to the version of the dataset hosted on Kaggle.

**Correction of AdaptableSnail data sampling rate errors.** A subset of AdaptableSnail videos were acquired at 25 fps rather than 30 fps. In the original competition release, the behavior annotations for these videos were resampled at 30 fps but the pose was not, leading to a mismatch between pose and behavior. This was corrected by regenerating annotation files without resampling, so both annotation and pose for affected videos are at 25 fps.

**Correction of AdaptableSnail mouse ID swaps.** A small subset of AdaptableSnail videos showed low classifier performance for multiple behaviors. Visual inspection revealed that these videos contained segments of time where the identity of two animals were swapped. Leveraging the fact that mice in these videos have dyed coats, we detected and corrected ID swaps in affected videos using an image color filter on the raw videos.

**Correction of AdaptableSnail and InvincibleJellyfish list of annotated behaviors.** Five AdaptableSnail videos were indicated as having been annotated for approach, submit, and rear, however these behaviors had not been scored. Additionally, 33 InvincibleJellyfish videos were incorrectly indicated as having been annotated for allogrooming. These behaviors were removed from the “behaviors labeled” field in the metadata csv file.

**Removal of InvincibleJellyfish escape behavior.** This behavior was identified as having unusually low classifier accuracy compared to other labs’ escape classifiers. Visual inspection of the videos alongside manual annotations revealed that the escape label was applied to two distinct behaviors: when an intruder animal escaped aggression from the resident, and when an animal jumped upwards attempting to escape the home cage, a behavior that often occurred spontaneously without resident aggression. The label was therefore removed from the dataset.

**Correction of InvincibleJellyfish time offset error.** A subset of videos in this dataset showed a fixed-per-video 0.5-2 second offset between pose+video and behavior annotations. This was corrected by applying a video-level shift in annotation timing, using the dataset’s manual sniff annotations and a pose-based feature detecting proximity of the resident’s nose to the intruder.

**Correction of LyricalHare approach behavior.** This behavior was identified as having unusually low classifier accuracy compared to other labs’ approach classifiers. Visual inspection of the videos revealed that the subject->target directionality of the approach label had been reversed. This was corrected in the revised dataset.

**Rebuild of LyricalHare, JovialSwallow, and InvincibleJellyfish annotation files.** A small number (<20) of annotation files were generated incorrectly or paired with the wrong pose file. These were identified by screening for videos with unusually low classifier accuracy, and were rebuilt from source data.

**Removal of PleasantMeerkat attack/chase/escape behaviors.** These behaviors were identified as having unusually low classifier accuracy compared to the same behaviors scored by other labs. Visual inspection revealed that all annotations for these actions lasted for a single frame, and denoted the onset of the behavior rather than a frame-wise label of its occurrence. The behavior labels were therefore removed from the dataset.

**Correction of SparklingTapir and DeliriousFly misreported framerates.** A subset of videos in the SparklingTapir and DeliriousFly were acquired at high framerate and downsampled prior to pose estimation and annotation; this downsampling had not been accounted for in the reported framerate. We resampled the source videos to match the framerate of the pose and annotation data, and framerates adjusted in the metadata.

**Creation of a "sniff-all" label.** We found during post-competition evaluation that the "sniff" classifier showed high variation between labs, due in part to differences in which behaviors were included in the sniff category. Whereas some labs annotated all social investigation behaviors as "sniff", other groups separated particular types of sniffing (such as face or anogenital-directed) into a separate category, resulting in a sniff classifier that excluded those types of action. We therefore added another classification head for "sniff-all", which was trained and evaluated by merging all sniff annotation categories (recipriocalsniff, sniffbody, sniffgenital, sniffface, and sniff) for any lab where both sniff and another subtype of sniffing were annotated: affected labs were CautiousGiraffe, GroovyShrew, InvincibleJellyfish, NiftyGoldfinch, and TranquilPanther. For consistency with other labs, we used the sniff-all classifier in place of the lab's original sniff classifier in all evaluations in this paper.

#### HIDRA classifier architecture, training, and inference

The HIDRA model is adapted from the first-place Kaggle solution described above, with a minor modification to model architecture. Specifically, the first-place solution used weight sharing between classifiers for the same-named behavior from different labs, raising the possibility that poor performance on data from some labs could be due to either overfitting to other labs, or lab-to-lab differences in how behaviors are defined. We therefore extended the supervised classification network to use a distinct classification head for each of the 82 lab-behavior pairs; the shared heads from the original architecture were still preserved, but only used during training. This modestly improved or at least failed to degrade classifier performance across all tested behaviors.

**HiDRA training.** Training of HiDRA follows the approach of the first-place Kaggle solution. We train the self-supervised feature extractor and supervised classifier components of HiDRA separately. Each model stage processes an overlapping temporal window of 64 sampled frames, with an additional 32 frames of temporal context padded on each side. During training of both stages, each sampled window is augmented with independent random transformations, including random rotation, random reflection/flipping, random isometric scaling, random temporal dilation, and additive Gaussian keypoint jitter; scaling, dilation, and jitter bounds are provided in Table 4.

First, we trained the feature extractor in a self-supervised manner to forecast the displacement of each keypoint on the next frame, quantized into 16 radial magnitude and 16 angular bins, and trained with a cross-entropy loss with masking over frames and keypoints that are missing, and at sequence boundaries. We ran training for up to 125,000 steps using the Adam optimizer with a learning rate of 0.02, batch size 128, early stopping (patience 15,000 steps), and exponential moving averaging of weights (decay 0.9993), repeating training for each of the five model configurations outlined in the first place solution.

We next freeze the weights of the self-supervised model, and connect its output to the supervised classification architecture (feature merging, temporal filtering, and classification heads). We train the supervised architecture with a per-frame sigmoid binary cross-entropy loss, masked to restrict training to only the behaviors annotated within each training video. We ran training for up to 50,000 steps using the Adam optimizer with learning rate 0.004, batch size 128, early stopping on a 15% validation split of the training set (patience 10,000 steps), and exponential moving averaging of weights (decay 0.9993).

Classifier head outputs are probability values, which we convert to a binary label with a per-head decision threshold. We calibrate the threshold on the training dataset by sweeping over a range of threshold values and selecting the value that maximizes that head's F1 score on the training dataset.

**HiDRA inference.** Inference with HiDRA also follows the approach of the first-place Kaggle solution. During inference, the model combines three averaging methods to compute the probability of each behavior label. Because each sample fed to the model is a 64-frame sequence, we can perform prediction on neighboring, overlapping windows. We also perform random test-time augmentation (random rotation, flip, scale, and time dilation), and average all predicted probabilities for each window and augmentation configuration that covers each frame. Finally, because each model in the ensemble resamples the behavior data to a different framerate (see Table 4), we resample each probability series to the video's native resolution, and perform averaging over ensemble members. We apply the fit decision threshold to this final average to produce binary behavior labels.

**Frame-wise aggregation of HiDRA labels.** By default, HiDRA's 82 classifier heads can produce multiple positive labels per frame: this may include multiple labs' classifiers firing for the same behavior (eg attack), or multiple classifiers firing for different but related behaviors that are scored differently by different labs (eg both "sniffface" and "reciprocalsniff" firing during reciprocal sniffing).

These labels may be used as-is, however in some cases it may be desirable to combine them to produce a single behavior label per frame, for example to override the output of low-performing classifier heads when a more high-performing head also fires. We use this approach in our analysis of the autism and multi-day experience datasets, with some exceptions described here. A naive approach to achieve a single behavior label per frame is to select whichever classifier head's probability output is the highest above that classifier's threshold. However, this potentially leads to underestimating the frequencies of behaviors that co-occur such as sniffface and reciprocalsniff. We therefore added ontology-based exceptions to the winner-take-all rule; these are classes of behavior labels that are permitted to co-label frames:

1) Sniffing subtypes: sniff, sniffbody, sniffface, sniffgenital, and reciprocalsniff can all be co-emitted.
2) Mounting/dominance: attemptedmount, mount and intromission can be emitted alongside dominance or dominancegroom.

This list can be extended based on prior knowledge of which behaviors are likely to co-occur.

#### Comparison to video-based feature extractor

To evaluate the efficacy of video-derived features for behavior classification in the DooM benchmark, we adapted the HiDRA architecture to ingest spatiotemporal representations extracted using VideoPrism on the raw videos, substituting this for the self-supervised GNN element of HiDRA (Fig 2A). This enables us to compare representations extracted from a frozen video backbone pre-trained on general Internet data against the representations learned by the trajectory model using mouse-specific data.

**Video feature extraction.** We use the public pretrained VideoPrism encoder(*4*) (`videoprism_public_v1_base`(*4, 5*)) with frozen weights to extract features directly from video frames. Each video was rescaled (aspect-ratio-preserving) such that the entire image fit within 288 × 288 pixels, with the remaining pixels zero-padded. Input videos were parsed into 16-frame clips sampled at a temporal stride of 8 frames for computational efficiency. The frozen encoder processed each 16-frame clip into a tensor of dimension 16 frames × 256 spatial patches × 768 feature dimensions (256 × 768 feature map per timestep). To reconstruct continuous per-frame visual features from overlapping clips extracted at a temporal stride of 8 frames, we use slices from adjacent clips: for the initial clip of a sequence, frames [0, 12) were retained; for all interior clips, frames [4, 12) were retained; and for the final clip, frames [4, 16) were retained.

**Model architecture.** The video classifier uses the same downstream classifier of HiDRA after feature extraction. The difference is that whereas the pose-based HiDRA model sums representations across body parts, the video model uses single-head attention pooling across the 256 spatial patches at each timestep (employing a single head to match the unweighted sum in HiDRA as closely as possible) to yield a single 192-dimensional feature vector per frame, aligning with HiDRA's node dimension after pooling. Similar to HiDRA, a learned laboratory identification embedding was weighted and added to this per-frame representation. The combined representations were subsequently passed through the same classifier architecture as HiDRA.

**Training and inference.** The video-based classification network was trained on the compiled training videos across all 15 DooM laboratories plus the CRIM13 dataset. CalMS21 features were not extracted for supervised classification due to the dataset's size (extracted features were 7 Tb for DooM and CRIM13 alone.) Classifiers were trained using masked binary cross-entropy loss and optimized using the Adam optimizer with a base learning rate of 0.004, a batch size of 128 windows, and an exponential moving average with a decay rate of 0.9993. Models were trained for a maximum of 50,000 optimization steps, evaluated every 2,000 steps, with early stopping triggered if validation F1 failed to improve over a patience window of 10,000 steps.

To compare against the ensemble of HiDRA, an ensemble of five model instances was trained under this scheme. Due to the high computational overhead of video feature extraction and the absence of direct video analogues for keypoint-specific geometric transformations (such as individual limb coordinate jitter or skeleton swapping), video ensemble members were trained without those pose-specific augmentations, retaining only HiDRA's temporal augmentation. All five ensemble members converged successfully to early stopping without premature preemption. For inference, test-set predictions were generated by evaluating sliding clip windows with the frozen encoder, assembling continuous per-frame features via temporal slicing, and passing 128-step windows through the five ensemble models, averaging predictions across windows and members.

### HiDRA performance analysis

#### Error decomposition and clustering

For each HiDRA classifier, we identified all errors on that lab's test set, and decomposed these errors into four categories:

- **Jitter errors:** false positive or false negative errors that occur within 0.5 seconds of a ground-truth-positive frame.
- **Missed bouts:** a ground-truth bout in which fewer than 30% of frames were detected by the classifier.
- **Misidentified:** a false positive output of the classifier that is not contiguous with a ground-truth bout and does not fall within 0.5 seconds of a ground-truth bout, but coincides with ground-truth annotations for a different behavior.
- **Extra bouts:** predicted bouts that do not fall within 0.5 seconds of a ground-truth bout and do not coincide with annotations for other behaviors. Includes frames within bouts that are contiguous with ground-truth but extend more than 0.5 seconds outside the duration of the ground-truth annotation.

We also investigated further subdividing of error types, but restricted our visualization to the above four for simplicity. In particular, jitter errors can be more finely dissected: while the jitter error category as defined above was dominated by differences in timing of bout onset and termination, it also includes brief gaps in the human ground truth annotations that were missed by the classifier, or brief exclusions by the classifier that were not excluded in the human ground truth annotations. Our jitter definition also does not distinguish between jitter errors arising when a behavior is immediately bordered by another label type (for example approach transitioning directly into investigation) vs errors arising in isolation from other behaviors. Including these error types did not significantly alter our conclusions about the classifiers.

For visualization, we computed the percent breakdown of each of the four error types defined above for each classifier, and clustered classifiers using agglomerative hierarchical clustering with Ward linkage and a Euclidean distance metric.

#### Within-lab F1 score prediction

To identify the factors driving classifier F1 score, we performed correlation and regression analysis relating F1 score to experimental metadata. We defined the following nine features as candidate predictors of F1 score:

- **Number of positive training frames:** the total number of frames annotated as the target behavior in the lab's training set, log-transformed.
- **Number of training bouts:** the count of discrete annotated bouts of the target behavior across the lab's training videos, where a bout is one contiguous annotated interval for a given agent–target pair. Log-transformed.
- **Mean bout duration:** the average duration (in seconds) of the annotated bouts of that behavior, log-transformed.
- **Behavior rarity:** we defined annotation prevalence as the fraction of scored frames that were positive for the behavior, and rarity as −log(prevalence) so that rarer behaviors have higher values.

- **Frame rate:** video acquisition rate in frames/second. For labs that had variation in frame rates across videos, we took the mode.
- **Number of tracked keypoints:** the number of body parts each lab tracked (see **Suppl Fig 1**.)
- **Video resolution:** spatial resolution of the lab's videos in pixels per centimeter. For labs that had variation in resolution across videos, we took the mode.
- **Lab identity:** a categorical variable identifying the recording lab, one-hot encoded.
- **Behavior identity:** a categorical variable identifying the behavior, one-hot encoded.

**Correlation analysis.** For numeric predictors, we computed the correlation between each predictor and F1 using Spearman's rank correlation coefficient (Spearman's $\rho$). For unordered categorical predictors, we instead used the correlation ratio $\eta$, defined as the variance of F1 scores within a category divided by the total variance across categories.

**Permutation importance test.** Because some of our regressors may be correlated with each other, we identified the features most strongly predictive of F1 score using a permutation importance test. Specifically, we trained a random-forest regression model to predict F1 score from predictors, where the values of randomly selected predictors were permuted across observations. For each predictor, we then computed the mean decrease in $R^2$ on held-out data when that predictor's values were permuted, averaged over 10 permutations per fold and over 40 cross-validation splits.

**Additional correlates of F1 score.** In addition to the features used in the above analysis, we note two features of classifiers themselves that we found to be correlated with F1 score. The first of these is the model threshold, used to binarize raw classifier output (probabilities) into discrete behavior labels; classifiers that fail to separate positive from negative frames have lower best-fit thresholds, with a Spearman's $\rho$=0.736 between threshold value and F1 score. A second strong correlate of F1 score is ensemble agreement. Following the first-place Kaggle solution, HiDRA uses an ensemble of five models trained with different amounts of temporal and keypoint downsampling. The mean pairwise Pearson's correlation between ensemble models was itself highly correlated with F1 score (Spearman's $\rho$=0.80).

Strikingly, several low-F1 score classifiers still had high ensemble agreement. Upon visual inspection, high-agreement false positives often corresponded to genuine instances of behavior that were missed by human annotators, for example, for NiftyGoldfinch - run and LyricalHare - approach. These are short, common behaviors that require an annotator to consistently apply internal scoring rules, for example, deciding how fast a mouse must move to be considered running-- a task that is difficult to perform, particularly across days or weeks of scoring. Inter-classifier agreement could therefore provide a means of testing for and correcting human-generated noise and drift in manually annotated datasets.

#### Cross-lab F1 score analysis

For the cross-lab F1 score analysis, we identified all behaviors that were annotated by more than one lab. For each behavior, we evaluated each valid classifier on the test set videos of each lab that scored that behavior. Because our objective was to measure similarity of behavior definitions rather than to quantify pure transfer accuracy, we permitted each classifier threshold to be tuned on each evaluation lab's training set, and report F1 score on the evaluation lab's test set. We note that the HiDRA pipeline includes a lab ID embedding: for all cross-lab experiments, we use the lab ID embedding of the lab that provided the classifier, rather than the ID of the lab that classifier is being evaluated on.

---

### Cross-lab performance experiments

#### Application to CalMS21

We evaluated HiDRA on the standard CalMS21 Task 1 test set, which consists of manual annotations of sniffing, attack, and mount behaviors. For each CalMS21 test behavior, we identified all DooM labs that annotated for that behavior and applied those labs' classifiers on the CalMS21 test set, preserving the lab ID embedding, classifier head, and classifier threshold of the source lab. Because the mount annotation in CalMS21 includes both mounting and intromission, we used classifiers from two labs that had annotated both behaviors, and merged each lab's mount and intromission predictions to create the final mount prediction. We computed the CalMS21 F1 score as described above.

We note that CalMS21 pose trajectories were included in the training data for the self-supervised embedding models of HiDRA. In these models, pose data alone is used to train a network to predict the pose on the next frame of a video given the pose history, and these weights are frozen prior to training of the supervised classifiers in HiDRA. During HiDRA's supervised training process, neither pose nor annotation data from CalMS21 was used.

#### Leave-one-lab-out experiment

To study the ability of HiDRA classifiers to generalize to unseen labs, we performed a leave-on-lab-out (LOLO) experiment. We selected five labs whose data spanned a range of appearances: AdaptableSnail, GroovyShrew, InvincibleJellyfish, LyricalHare, and NiftyGoldfinch. For each lab, we first trained a new version of the HiDRA supervised classifier model (feature merging, temporal filtering, and classifier head components, **Fig 2A**) with that lab's data withheld. In a test experiment with LyricalHare, we found that also excluding that lab's data from training of the self-supervised model did not substantially alter our results (data not shown)- therefore to reduce training time, the self-supervised model was not retrained, and only one of the five ensemble models was used (27fps_6bp).

For each held-out lab, we identified four to six social behaviors for testing, spanning a range of training dataset sizes. Four of these are behaviors unique to the held-out lab, for which no classifier head exists in the retrained HiDRA model. These are indicated by a star.

| | |
|---|---|
| AdaptableSnail: | chase, attack, avoid*, approach |
| GroovyShrew: | sniff, sniffgenital, approach, defend |
| InvincibleJellyfish: | attack, dominancegroom*, sniff, allogroom |
| LyricalHare: | attack, defend, sniff, approach |
| NiftyGoldfinch: | chase, follow, sniff, tussle*, sniffface*, sniffgenital |

We first computed F1 score of the re-trained (LOLO) HiDRA model for each held-out behavior prior to any fine-tuning, excluding behaviors unique to the held-out lab. For each held-out behavior, we identified the set of "donor" labs that annotated for that behavior, and ran inference on the held-out lab's test set using that lab's lab ID embedding, classifier head, and classifier threshold, reporting the F1 score of the best-performing donor head normalized by the F1 score of the full HiDRA model (**Suppl Fig 5A**). We found that tuning the threshold of the selected classifier often improved performance, therefore we also report the normalized F1 score of the model after tuning only the donor lab threshold, for which we used the held-out lab's training set (**Suppl Fig 5A**, red bars).

We next performed a fine-tuning experiment for each of the 22 held-out behaviors, sweeping over a range of training set sizes. For each held-out behavior, we first built a log-spaced grid of target positive-training-frame counts, with a floor of 100 frames and a ceiling of the full annotation set for that behavior. For each target, we drew a contiguous segment of frames from the training data, with video order randomly permuted and with training repeated over three such permutations per grid point.

In each grid point replicate, we randomly selected one donor lab that annotated for the target behavior, and initialized the model using that lab's embedding and classifier head. For behaviors with no donor head, lab ID embedding and head were initialized randomly. Fine-tuning trained the lab embedding, temporal filtering layers, and target behavior's classification head, equivalent to the "head + trunk mode" described in the held-out dataset experiment below. We trained the model on each held-out behavior separately, using the AdamW optimizer (8,000 steps, max learning rate 0.004 with cosine learning rate decay to 0 at 15,000 steps (unreached), exponential moving average weight decay with alpha=0.9993, and early stopping) with a binary cross-entropy loss on the target behavior. After training, we selected the classifier threshold via grid search to maximize F1 score on the full training set, then reported the F1 score evaluated on the test set, averaged over the three replicates.

#### Fine-tuning on a held-out dataset

##### Dataset

To evaluate HiDRA on a completely new dataset that it never saw during training, we collected a small dataset of home-cage behavior from a mouse model of amyotrophic lateral sclerosis (ALS). We recorded three cages, each housing three co-housed male hemizygous G93A SOD1 mice from The Jackson Laboratory (#004435), at 7–20 weeks of age, a period spanning the pre-symptomatic to symptomatic stages of neuromuscular degeneration. We monitored the animals daily for disease progression and euthanized them upon reaching humane endpoints (loss of the righting reflex within 30 s, or body weight below 80% of baseline). The Salk Institute for Biological Studies Institutional Animal Care and Use Committee approved all procedures (protocol 16-00011), and we performed them in accordance with the NIH Guide for the Care and Use of Laboratory Animals.

We acquired video at 50 FPS, with a 1280 × 1024 px image size and 35 px/cm resolution using a Basler acA1300-75gm camera (1280x1024, 88 fps, mono Python 1300), Tamron 12VM412ASIR 4-12mm F/1.2 IR (Day/Night) lens, and LP800-S40 Near-IR Longpass Filter, slip-mount, 40mm. We did not use any of these recordings at any stage of HiDRA's development. We estimated the pose of the mice using SLEAP, tracking 15 keypoints: the nose, ears, head, neck, shoulders, body center, hips, tail base, tail tip and three intermediate tail points. HiDRA uses seven of these: the nose, ears, neck, body center, tail base and tail tip. We manually proofread the SLEAP tracks to correct identity swaps and converted them to HiDRA's input format. A single trained annotator labeled three behaviors for every animal in six five-minute clips: self-grooming ("selfgroom"), anogenital sniffing ("sniffgenital") and jumping from a covered feeder cage down to the home cage bottom ("jumpdown"). We split the clips into a two-clip training set (24 selfgroom, 35 sniffgenital and 49 jumpdown bouts) and a four-clip evaluation set (37 selfgroom, 61 sniffgenital and 66 jumpdown bouts). We split by clip rather than by animal. The two training clips came from two of the three cages. The evaluation set contained one clip from each of those cages, recorded 6–8 weeks before the training clips, and two clips from the third cage, whose animals do not appear in the training set. Fine-tuning improved F1 as much or more on the two clips from this unseen cage as on the other two (mean F1 gain over zero-shot, unseen vs seen cages: jumpdown +0.72 vs +0.58, selfgroom +0.13 vs −0.01, sniffgenital +0.12 vs +0.06).

##### Annotation protocol

A single annotator labeled the same six clips under two protocols (fully manual and human-in-the-loop) using a custom-built annotation tool. In the first, manual protocol, the annotator searched each clip and drew every bout by hand (326 drawing actions, 272 bouts). The annotator completed the manual protocol in full, and later started the second, human-in-the-loop (HITL) protocol. The annotator did not see model output during the manual pass, so the manual labels are an independent reference. Because the HITL pass came second, however, familiarity with the clips may have shortened review, and the effort reductions we report may be optimistic.

##### Fine-tuning experiments

We fine-tuned each of HiDRA's five ensemble models on the fully manual labels of the two-clip training set described above, comparing two approaches:

- **Head-only mode** trains only the target behavior's head in the linear output layer: 256 readout weights, a gain, and a bias, or 258 parameters per behavior.
- **Head + trunk mode** also trains the lab embedding and temporal filtering layers (three BiLSTM + feed-forward blocks), as in the LOLO experiment; about 4.8M parameters per behavior.

Several DooM labs annotated selfgroom and sniffgenital, but none annotated jumpdown. We initialized the selfgroom and sniffgenital classifiers from the NiftyGoldfinch lab ID embedding and classifier heads for those behaviors. We initialized the jumpdown classifier from the NiftyGoldfinch lab ID embedding and climb classifier head, which achieved an F1 score of 0.07 on jumpdown before fine-tuning.

For each classifier and training mode, we trained the model with Adam (learning rate 0.004, batch size 128 windows, no weight decay) and a binary cross-entropy loss on the target behavior only. As in HiDRA training, we kept an exponential moving average of the weights (decay 0.9993) and used the averaged weights from the final step. We kept the input normalization statistics at their pretrained values. With these statistics fixed, head-only fine-tuning leaves every other classifier in the model unchanged, so a head-only checkpoint can replace the published model. Head + trunk fine-tuning instead changes the temporal filtering layers that all of HiDRA's classifier heads share. This alters the output of every head that we did not retrain, so we used each head + trunk checkpoint only to predict its own target behavior.

We trained the head-only models for 300, 1,000, 8,000, 16,000 or 32,000 steps without early stopping; the fine-tuned classifiers shown in **Fig 3l** use 8,000 steps. For selfgroom and sniffgenital, we used a cosine learning rate decay that reached zero at the final step. For jumpdown, we used a constant learning rate, which converged in fewer steps for this behavior.

We trained the head + trunk models for 1,000 steps at an effectively constant learning rate. For consistency with the LOLO experiment, we kept its 15,000-step cosine schedule, under which the learning rate is still 98.9% of its peak at step 1,000. A head + trunk step costs about 30 times as much as a head-only step (about 1 h per behavior for 1,000 steps), so we did not train these models longer.

We repeated each head-only condition with three random seeds, which varied the data order and augmentation; the warm start fixes the initial weights. Because we observed minimal variation across seeds in head-only mode, we trained the head + trunk models once per behavior. We report training times as the wall-clock time to train all five ensemble members sequentially on one NVIDIA RTX A6000 GPU.

Fine-tuning changes the scale of the classifier head's output probabilities, so we re-selected the decision threshold after training in both modes. We used a leave-one-clip-out procedure. For each test clip, we chose the threshold that maximized the F1 score on the other three test clips concatenated, found by grid search, and applied it to the held-out clip. As in our other analyses, we pooled frames before scoring: across all animals for the self-directed behaviors (selfgroom and jumpdown), and across all ordered subject-target pairs for sniffgenital. We report the F1 score averaged over the four test clips.

#### Human-in-the-loop experiments

To evaluate HiDRA as a tool for human-in-the-loop (HITL) annotation, we compared annotation time usage in pure manual vs classifier-assisted labeling. HITL annotation consists of using a classifier to propose a set of candidate behavior annotations; the annotator then reviews each of these, adjusting or discarding bouts when necessary. This approach can accelerate annotation particularly of rare behaviors, because it allows annotators to skip inspection of large portions of the video. Because the review process only involves inspection of candidate bouts, and not of negative regions, HITL tools typically set a low threshold for candidate bout detection, excluding from inspection only segments where the model is confident that the behavior did not occur, at the cost of producing a higher proportion of false positives.

In testing our classifier-assisted annotation workflow, a human annotator (“user”) working within our custom annotation interface ([https://github.com/talmolab/laras-labeler](https://github.com/talmolab/laras-labeler)) first used HiDRA to generate per-frame probabilities of the selfgroom and sniffgenital behaviors on the four test video clips (done by omitting the final thresholding step of HiDRA), using the published NiftyGoldfinch HiDRA classifiers with no fine-tuning. Because HiDRA has no jumpdown classifier, for this behavior the user instead manually labeled 49 bouts in our two training clips, performed a single round of fine-tuning using the NiftyGoldfinch climb classifier head in head + trunk mode, then used this fine-tuned model to generate framewise probabilities of the behavior on our four test clips.

The user next reviewed HiDRA-generated outputs on the four test clips. Candidate bouts were frames above a single threshold, fixed before annotation and not changed (0.55 for selfgroom and jumpdown, 0.45 for sniffgenital). The user reviewed every candidate until the queue for each animal was empty, accepting (possibly with boundary edits), merging, or rejecting each. Round 1 of HITL annotation produced 1,790 decisions: 331 accepted, 69 merged and 1,390 rejected. Among accepted proposals, the user moved the start later in 215 cases and the end earlier in 252 (median start +10 frames, median end -26 frames). The median accepted proposal was 117 frames long, while the median retained bout was 46 frames. Accepting therefore took a median of 11.7 s, compared with 13.0 s to draw a bout by hand (drawing time only, excluding search). We then fine-tuned each classifier;s output head on the round-1 labels (head-only, 8,000 steps), predicted again, and the user reviewed the new candidates, which covered only frames still unlabeled after round 1 (round 2: 1,032 decisions, 69 kept). Proposals rejected in round 1 could not be proposed again.

Our annotation tool recorded the user's annotation time as an event log of actions, excluding idle spans over 120 seconds and counting any single pause as at most 15 s. We divided manual time into drawing (the summed duration of drawing actions) and searching (the rest of active time); HITL time is review time. We defined effort per labeled bout as each protocol's active time divided by the number of positive bouts it produced. The jumpdown seed bouts came from the manual pass, so we estimated their cost from manual rates. We prorated drawing time by bout count (49 of 115 bouts) and search time by footage (2 of 6 clips). We added this cost to HITL time and counted the seed bouts in HITL's output (63 reviewed + 49 seed = 112 bouts). Without the seeds, reviewing jumpdown took 26.4 s per bout, compared with 64.2 s per bout manually.

After both HITL rounds, bout yields differed between protocols (HITL vs manual): 77 vs 61 for selfgroom,206 vs 96 for sniffgenital, and 114 vs 115 for jumpdown. Round 2 found 2 selfgroom and 2 jumpdown bouts that round 1 had missed.HITL sniffgenital bouts were also shorter (median 0.87 vs 2.20 s). Of the 206 HITL sniffgenital bouts, 105 (median 0.42 s) overlapped no manual bout; some may be brief bouts missed in the manual pass. HITLalso missed bouts: 13 of the 61 manual selfgroom bouts and 1 of the 66 manual jumpdown bouts in the reviewed clips had no HITL counterpart, mostly because the annotator rejected the proposal (8 of 13 and 1 of 1). The experiment used one annotator and one pass per protocol, so there are no replicates.

#### Pose degradation analysis

To determine how HiDRA's performance depends on the quality of pose estimation, we performed a pose degradation analysis on the CalMS21 task 1 dataset. The poses released with CalMS21 were estimated with MARS (*6*), whose pose model was trained on manual annotations of seven keypoints (nose, ears, neck, hips and tail base) on 13,500 frames of video and evaluated on a 1,500-frame held-out test set. Some of the videos that supplied MARS's pose training frames are themselves part of the CalMS21 task 1 dataset.

We used these annotations (1024 × 570 px grayscale frames) to train new SLEAP pose models. We converted them to SLEAP format and randomly split the 15,000 frames into a 12,000-frame training partition, a 1,500-frame validation set and a 1,500-frame test set. From the training partition we drew subsets of 10, 50, 100, 500, 1,000 and 10,000 frames without replacement, using a single seeded random number generator, and we used the same validation and test sets for every subset.

For each subset we trained a two-stage top-down SLEAP model (sleap-nn v0.3.0), holding all other settings fixed. The first stage, a centroid model, located each animal by its neck keypoint. It used a UNet backbone (36 base filters, filter growth rate 1.5, max stride 32, output stride 2, confidence-map $\sigma$ = 2.5 px) on images downsampled by a factor of 2. The second stage, a centered-instance model, predicted all seven keypoints within a crop centered on the neck. It used a UNet backbone (24 base filters, filter growth rate 1.5, max stride 64, output stride 4, $\sigma$ = 1.5 px) on crops downsampled to two-thirds size. SLEAP set the crop size automatically from the largest animal in each training subset: 384 px for 10 frames, 448 px for 50–1,000 frames and 768 px for 10,000 frames. We trained both stages with random rotation (±180°) and scaling (0.9–1.1×) augmentation, using the Adam optimizer (learning rate $10^{-4}$) with a batch size of 8 split across two NVIDIA RTX A6000 GPUs. We halved the learning rate when validation loss plateaued and trained for up to 200 epochs of at least 200 steps each. We early-stopped the centered-instance models after 10 epochs without improvement and kept the checkpoint with the lowest validation loss. Centroid confidence maps are almost entirely background, so validation loss is a poor signal of detection quality when training data are scarce. We therefore trained every centroid model for the full 200 epochs without early stopping, saved a checkpoint at every epoch, and kept the one that detected the largest fraction of ground-truth animals in the validation set.

We then ran each trained model pair on every CalMS21 task 1 video, with a confidence map peak threshold of 0.2 and integral (subpixel) peak refinement. To measure pose quality separately from identity tracking, we assigned predicted instances to the two CalMS21 animal identities frame by frame, by Hungarian matching against the original CalMS21 poses on object keypoint similarity (OKS). We discarded matches with OKS < 0.3 and left unmatched animals as missing values. We exported the resulting tracks in the CalMS21 keypoint format and gave them to HiDRA unchanged.

For each trained model, we computed the 90% error radius of each keypoint, i.e. the 90th percentile of the Euclidean distance between predicted and annotated positions, on our 1,500-frame test set. To isolate keypoint localization from detection, we applied the centered-instance model to crops centered on the ground-truth animals, and we excluded keypoints that the model did not predict.

For each re-tracked version of CalMS21, we performed behavior classification using all HiDRA classifier heads for attack, sniff, and mount + intromission, as described above. **Figure 3M** reports the average classifier performance across heads.

---

### HiDRA applications

#### Test datasets

**Autism-model lines.** The autism line dataset analyzed here is described in Segalin et al(*6*). Briefly, this experiment tests group-housed male mice from five mouse lines (*Chd8*$^{+/-}$, *Cul3*$^{+/-}$, *Nlgn3*$^{+/-,}$ BTBR, and C57Bl/6J) in a standard resident-intruder assay. The Chd8, Cul3, and Nlgn3 lines are knock-out lines of genes associated with autism; animals were genotyped and cohorts of animals heterozygous for the gene knockout were compared to wild-type littermate controls. BTBR is an inbred line with low sociability, and was compared to C57Bl/6J animals as a control. Animals were tested at either 11-13 weeks of age (Nlgn3, BTBR, C57Bl/6J) or at 35-50 weeks of age (Cul3, Chd8) in a standard resident-intruder assay performed as in prior work(*7*–*9*). In the first, “group-housed” condition, experimental mice (“residents”) were tested in their homecage, with cagemates removed. Following a 5-15-minute habituation period, an unfamiliar group-housed male BALB/c mouse (“intruder”) is introduced to the cage, and animals are allowed to freely interact for a period of approximately 10 minutes. The resident male is then single-housed for two weeks, a manipulation that promotes aggressive behavior(*10*), after which a second resident-intruder test is performed in this “single-housed” condition. Both tests were recorded with infrared video captured at 30 fps from a top-view camera (Point Grey Grasshopper3), and pose estimation was performed using MARS(*6*). Sample sizes for all analyses are as follows:

| cohort | Chd8 | | Cul3 | | Nlgn3 | | BTBR | |
|---|---|---|---|---|---|---|---|---|
| geno | het | wt | het | wt | het | wt | BTBR | C57 |
| GH n | 8 | 8 | 8 | 7 | 10 | 9 | 10 | 10 |
| SH n | 8 | 8 | 8 | 7 | 8 | 8 | 10 | 10 |

**Multi-day experience.** The multi-day experience dataset analyzed here is described in Karigo et al(*9*). Prior to testing, 8- to 12-week-old C57Bl/6N male mice were individually cohoused with a BALB/c female mouse for one week, with the female mice removed from the cage one day prior to testing. On each day of testing, an unfamiliar hormone-primed BALB/c female mouse or intact BALB/c male mouse was introduced into the subject animal's home cage, and animals were allowed to freely interact for approximately ten minutes. Both tests were recorded with infrared video captured at 30 fps from a top-view camera (Point Grey Grasshopper3), animal pose was tracked using MARS, and videos were manually annotated frame-by-frame for investigation (sniffing), mounting, intromission, and attack. In the original dataset, ultrasonic vocalizations were also recorded, and used to identify individual mounting bouts as "USV+" or "USV–" based on the presence or absence of ultrasonic vocalizations during mounting; we do not make use of audio data here, and report only the presence of mounting.

#### Behavioral statistics

For both the autism and multi-day experience datasets, we ran all 82 HiDRA classification heads on each experimental file. We then merged the raw behavior annotation outputs of HiDRA using the protocol outlined under "**Frame-wise aggregation of HiDRA labels**" to produce annotation rasters for each animal.

**Genotype and housing contrasts.** To test for effects of genotype and housing on animal behavior, we first converted HiDRA's behavior annotation rasters to a single number per behavior, the fraction of time spent engaging in that behavior, by summing the number of positively annotated frames and dividing by the data frame rate. For each scored behavior, we tested for a significant difference in the fraction of time spent engaging in that behavior between heterozygotes and wild-type littermates within the group-housed condition (**Figure 4C**) using a two-sample t-test with Cohen's d (pooled standard deviation) as effect. We tested for an effect of housing condition on behavior in the heterozygotes of each strain (**Figure 4D**) using a paired t-test to compare the fraction of time spent in each behavior in the group-housed (GH) vs socially isolated (SI) condition.

To examine the effect of genotype on behavior bout durations, we computed the average bout duration for each behavior for each animal, where one bout is defined as a contiguous segment of video frames positively annotated for that behavior. For each scored behavior, we tested for a significant difference in the average bout duration (**Suppl Figure 9**) using a two-sample t-test with Cohen's d (pooled standard deviation) as effect.

In **Figure 4C-D**, a cell is marked with hatches if none of the animals in the groups being compared exhibited that behavior. In **Suppl Figure 9** (comparing bout durations), we require both groups to exhibit bouts of the behavior for bout duration to be quantified, therefore a cell is hatched for a behavior when either one or both sets of animals being compared did not exhibit that behavior. If all cells of a given figure column would be hatched, we omit that column from the figure.

**Behavioral transitions.** We examined the transition probabilities between HiDRA-annotated behaviors in each genotype and housing condition, selecting the single-housed Nlgn3 cohort for visualization in **Figure 4E** as the GH and SH conditions showed a large number of differences in computed transition probabilities. To construct the transition matrix, we first defined seven aggregate "states" consisting of the seven behavior categories defined in **Figure 1G**, as well as an eighth state corresponding to no annotated behavior. Using these eight behavior categories, we created a one-hot label of behavior state for each frame of video from each animal; to reduce transitions into the "no behavior" category, "no behavior" periods that were shorter than one second were omitted. For each animal we then constructed an empirical transition matrix by counting, for every pair of states A and B, how often a bout of state A was immediately followed by a bout of state B. We then divided by the number of times the animal left the state A to get the transition probability $p(B|A)$. Animals that never showed a certain behavioral category were excluded from the transition computation from that category.

To test for effects of genotype on behavior transition probabilities, we first defined the set of eligible transitions for testing as those for which the transition happened at least once, and in more than half of the animals compared, its row had at least five exits. Fig 4E compares single-housed Nlgn3 heterozygotes with wild types (8 versus 8) by t tests. Bars reflect every cell with uncorrected $p < 0.05$, as the heterozygote minus wild-type difference in mean probability (all 32 tested cells are in the source data); starred bars show significance after correction for multiple comparisons using Benjamini–Hochberg $q < 0.05$ over 32 tests.

**Nlgn3 post-attack behavior.** This analysis follows up the aggressive to appetitive transition of **Figure 4E**. An approach is almost always preceded by the two mice moving apart, so more approaches after an attack could mean either that heterozygotes separate more often or that they come back more often once separated. The HiDRA labels alone do not tell us which of the two has occurred, so we used the tracking to measure separation. We defined each mouse's position as the average of its tracked body points (here the nose, ears, neck, hips, and tail base). In the two seconds after each attack ended, we tested whether the mice moved apart more than 12 cm on any frame, and whether the resident subsequently approached the intruder (defined as HiDRA producing an approach label on any frame). We restricted this analysis to animals with at least five attack bouts, a cohort of 8 heterozygous and five wild-type animals.

**Multi-day analysis.** We cut every session to the length of the shortest session (556.3s) to compare equal recording times, then ran all HiDRA classifiers on the dataset and performed behavior label merging using the protocol outlined under "**Frame-wise aggregation of HiDRA labels**". We compared the time in each category between day 1 and day 3 within each intruder-sex group by paired t tests, with correction for multiple comparisons using Benjamini–Hochberg over the seven behavioral categories.

#### Comparison to MoSeq

**Decoding genotype.** We evaluated the ability of a trained classifier to distinguish heterozygous mice in the autism lines dataset from their wild-type littermates given the animals' behavior in the group-housed sessions. We compared three different sets of features: HiDRA bout statistics, HiDRA bout-conditioned features, and Keypoint-MoSeq.

*Bout statistics.* As a baseline, we used a summary of HiDRA's frame labels over the seven behavioral categories. This summary consists of the fraction of time annotated for each behavior, the average number of bouts per minute in each behavior, and the mean bout length of investigative, appetitive, and nonsocial behaviors (the three most common categories).

*HiDRA-conditioned features.* This more fine-grained representation consists of the resident's posture and movement within each behavior. The complete feature set consists of bout, posture, and movement features. First, we took the classifier head with highest-probability output on each frame, and for each behavior computed the share of frames on which that behavior's probability was the highest among all classifier heads. We next computed postural features for each of 12 behaviors, by taking the average of 14 distances between keypoints that describe animal posture (normalized by per-session body size) conditioned on the animal performing that behavior. Finally, we computed movement features for attack, dominance, freeze and sniffing behaviors, consisting of the 10th, 25th, 50th, 75th and 90th percentiles of speed and turning rate conditioned on that behavior occurring, split into a slow part (5-frame moving average) and a fast remainder.

*Keypoint-MoSeq*. The features are the fraction of each session's frames spent in each of K = 150 movement motifs (syllables), from models fitted to the resident's seven body keypoints on all autism sessions. We followed MoSeq's package protocol to fit the model: latent dimension 10, 3 lags, centroid noise estimated from the data, 50 iterations of autoregressive training and then 500 for the full model. We used self-transitioning priors $\kappa=10^8$ (autoregressive) and $4x10^6$ (full model) for a median syllable length of 400ms. Since MoSeq fits are quite variable, we fitted 20 models, and compared HiDRA to the MoSeq fit with median model performance over the 20 fits.

To decode genotype, we used each feature set as input to a logistic regression classifier with L2 regularization (C = 0.1), using feature standardization. We used leave one mouse out (LOMO) cross validation. The predictions are pooled into one area under the ROC curve (AUC) per line. We compute 95% bootstrap intervals for each AUC, by resampling mice with replacement, separately among mutants and controls and recomputing the AUC.

### Supplementary Figures

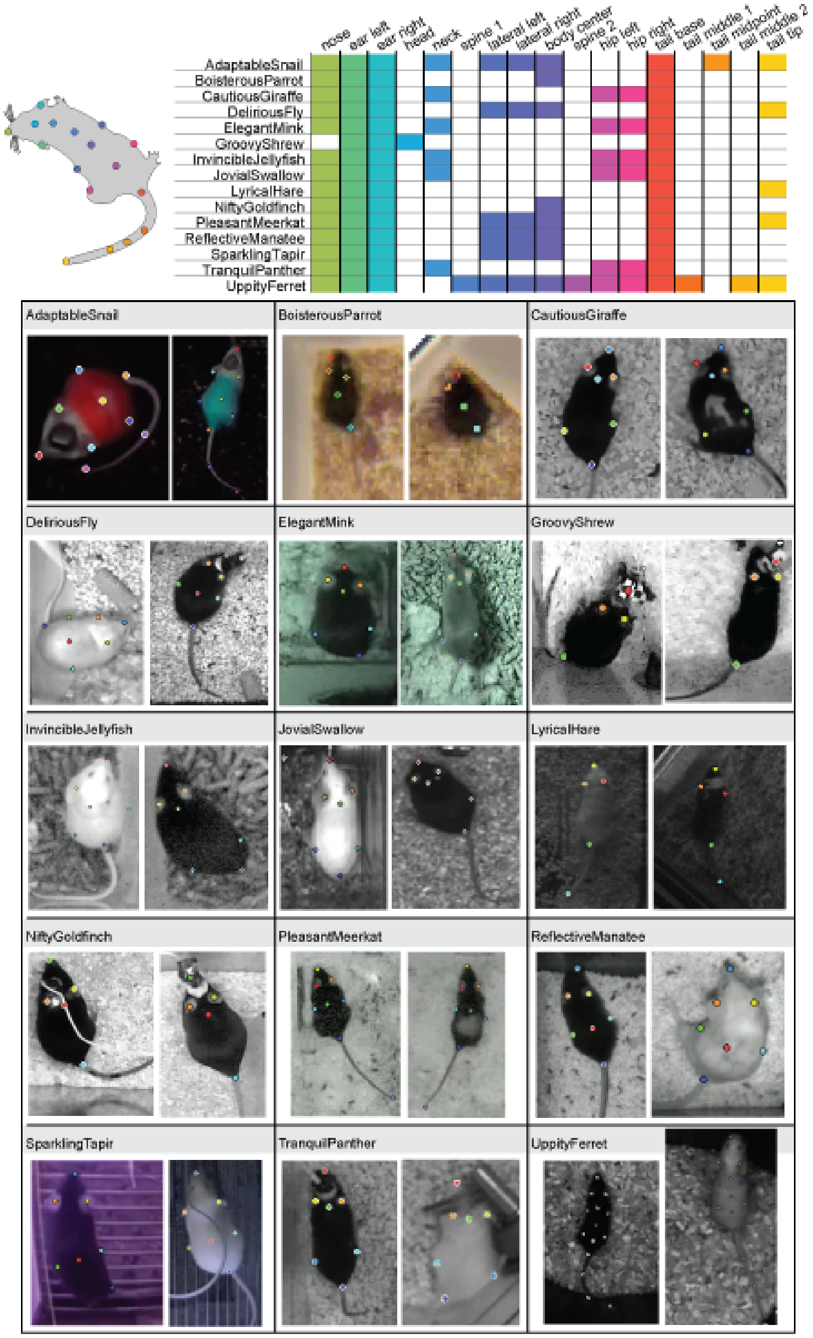

**Supplementary Figure 1. Pose annotations in the DooM dataset.** Above: Location of keypoints tracked across all labs in the dataset, and matrix showing keypoint tracking per lab. Not shown: in some AdaptableSnail videos, eight additional keypoints tracked the upper and lower corners of a rectangular device on the head of each animal. Below: representative images of mice from each dataset showing locations of tracked keypoints.

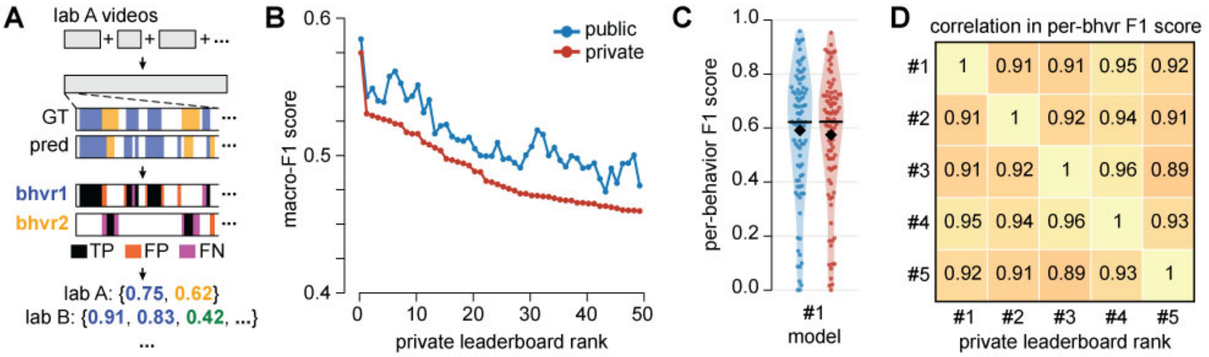

| | #1 | #2 | #3 | #4 | #5 |
|---|---|---|---|---|---|
| #1 | 1 | 0.91 | 0.91 | 0.95 | 0.92 |
| #2 | 0.91 | 1 | 0.92 | 0.94 | 0.91 |
| #3 | 0.91 | 0.92 | 1 | 0.96 | 0.89 |
| #4 | 0.95 | 0.94 | 0.96 | 1 | 0.93 |
| #5 | 0.92 | 0.91 | 0.89 | 0.93 | 1 |

**Supplementary Figure 2. Kaggle competition. A)** Procedure for computing macro-F1 score, details in Methods. **B)** Macro-F1 score on public and private test sets for top 50 submissions on the Kaggle competition leaderboard at the end of the two-month competition period. **C)** Distribution of F1 scores over all lab-behavior pairs in the public and private test set for the first-place Kaggle submission, showing high variance in classifier performance. **D)** Pearson's correlation between F1 scores over behaviors for the top five ranked submissions, showing that model performance over behaviors was highly consistent across top submission architectures.

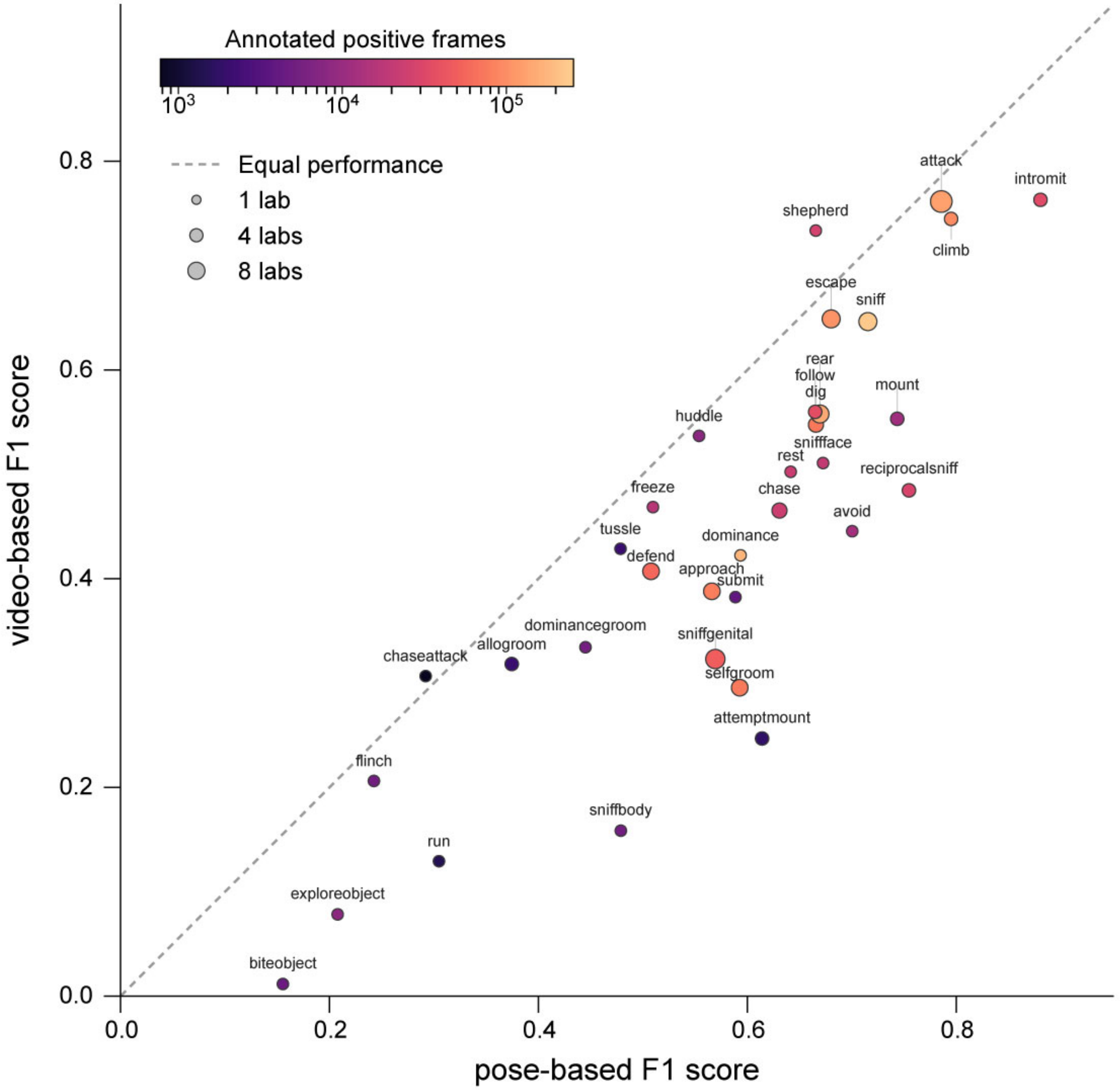

**Supplementary Figure 3. Video model analysis.** Performance of the HiDRA supervised classification trunk, trained and tested with either pose-derived features extracted using a self-supervised GNN, or video-derived features extracted using VideoPrism. The model trained using pose-derived features outperforms the video features for all but two~~one~~ behaviors (shepherd and chaseattack).

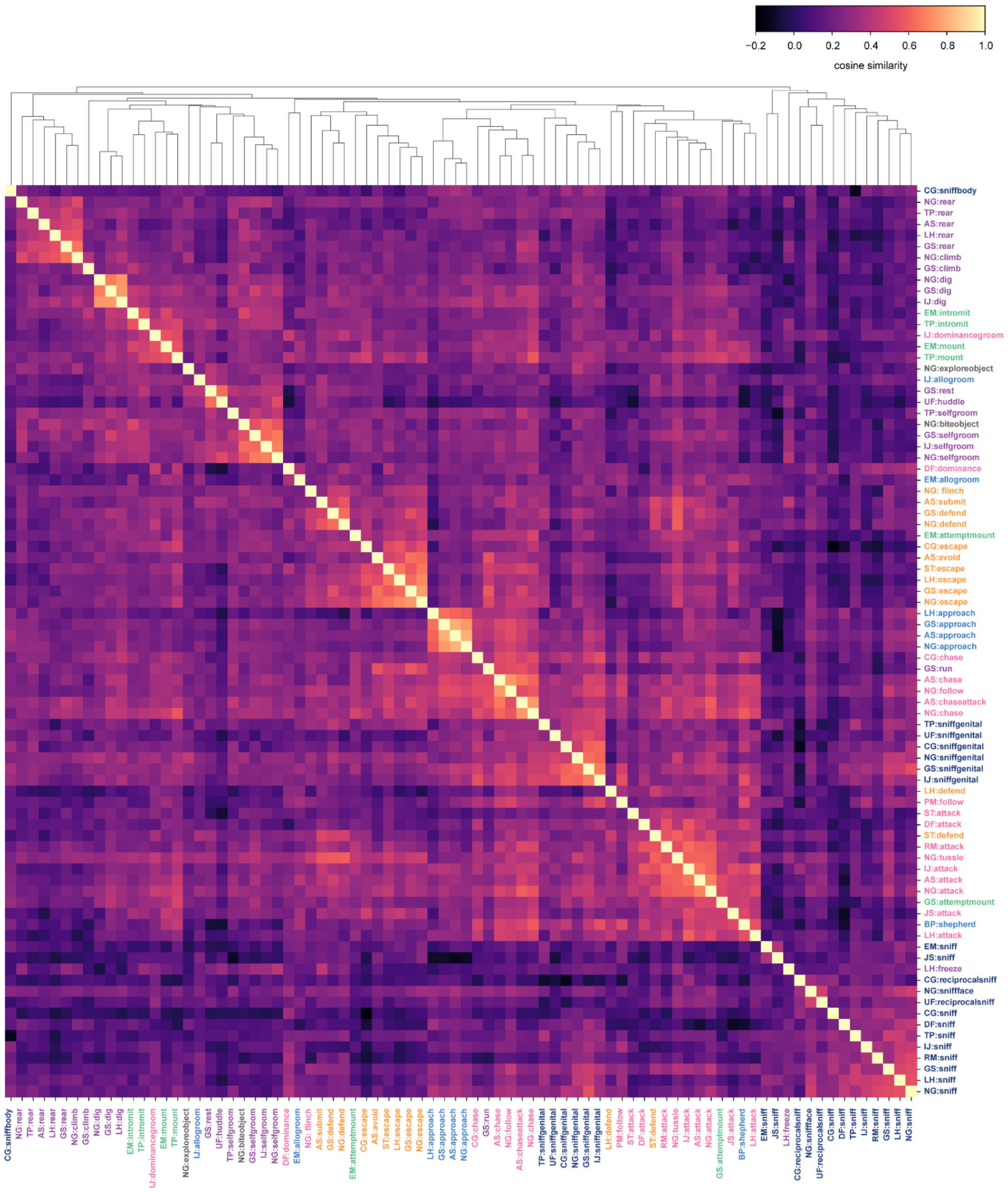

**Supplementary Figure 4. HiDRA classifier similarity analysis.** All HiDRA classifiers share a common trunk, with individual behavior classification performed by a single linear readout plus a head-specific scale and bias term (**Fig 2A**). Plotted is the cosine similarity of this 256-dimensional readout vector for each classifier from each lab, ordered by hierarchical clustering. Classifiers for the same behavior from different labs are often more similar to each other than classifiers for different behaviors within the same lab.

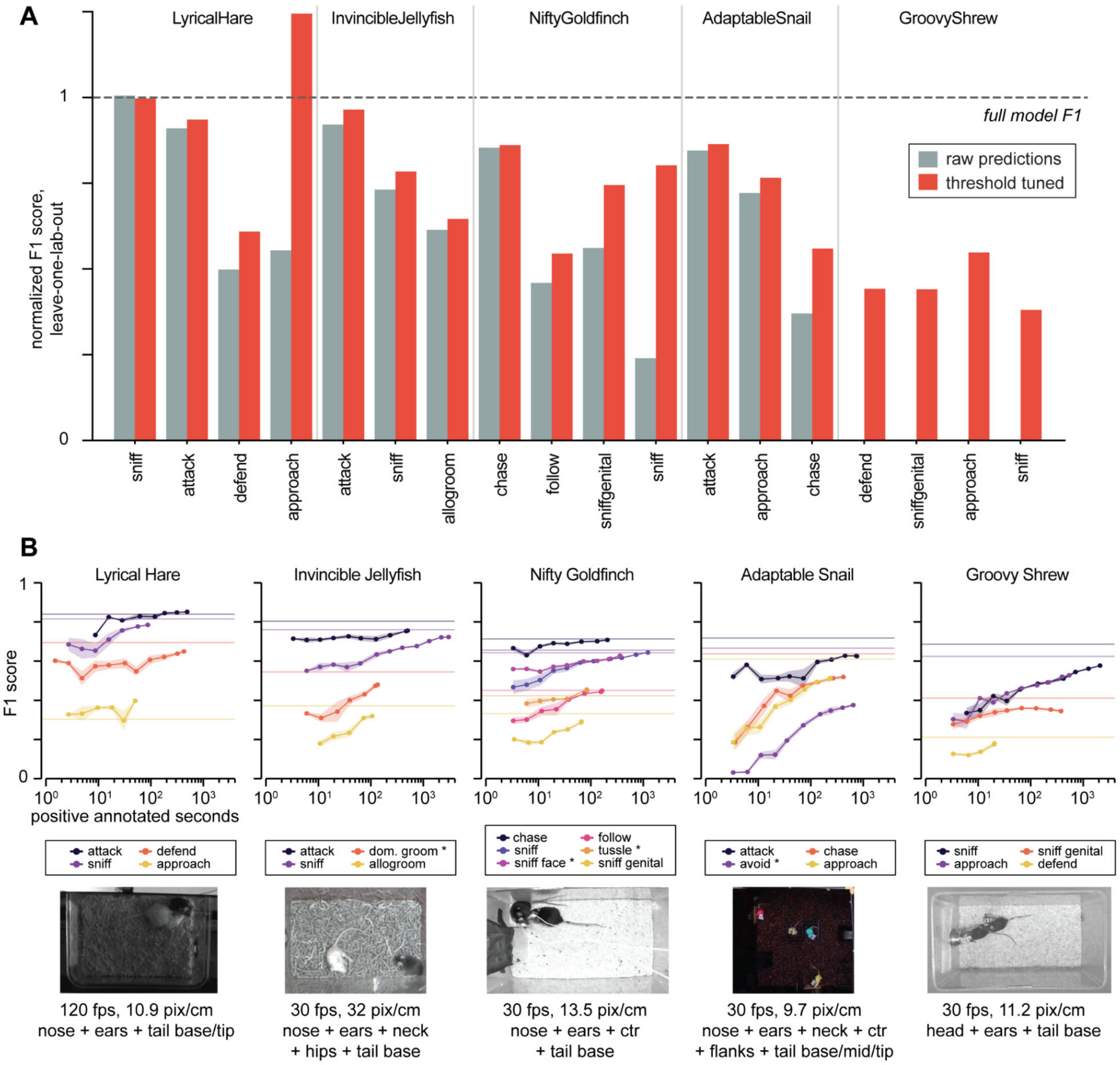

**Supplementary Figure 5. Leave-one-lab-out generalization. A)** F1 score of best-performing HiDRA classifier head for behaviors scored in held-out labs, evaluated on those labs' test sets. Gray bars use the existing HiDRA classifiers and decision thresholds; red bars reflect performance following tuning of the decision threshold (a scalar variable) on the training data from that lab. **B)** Fine-tuning of HiDRA classifiers on five held-out labs in the leave-one-lab-out experiment, showing F1 score as a function of training set size; horizontal lines indicate performance of baseline HiDRA model. Sample video frames, video metadata, and body parts tracked shown below each plot. * indicates behaviors not annotated by other labs (no warm-start head available for training.)

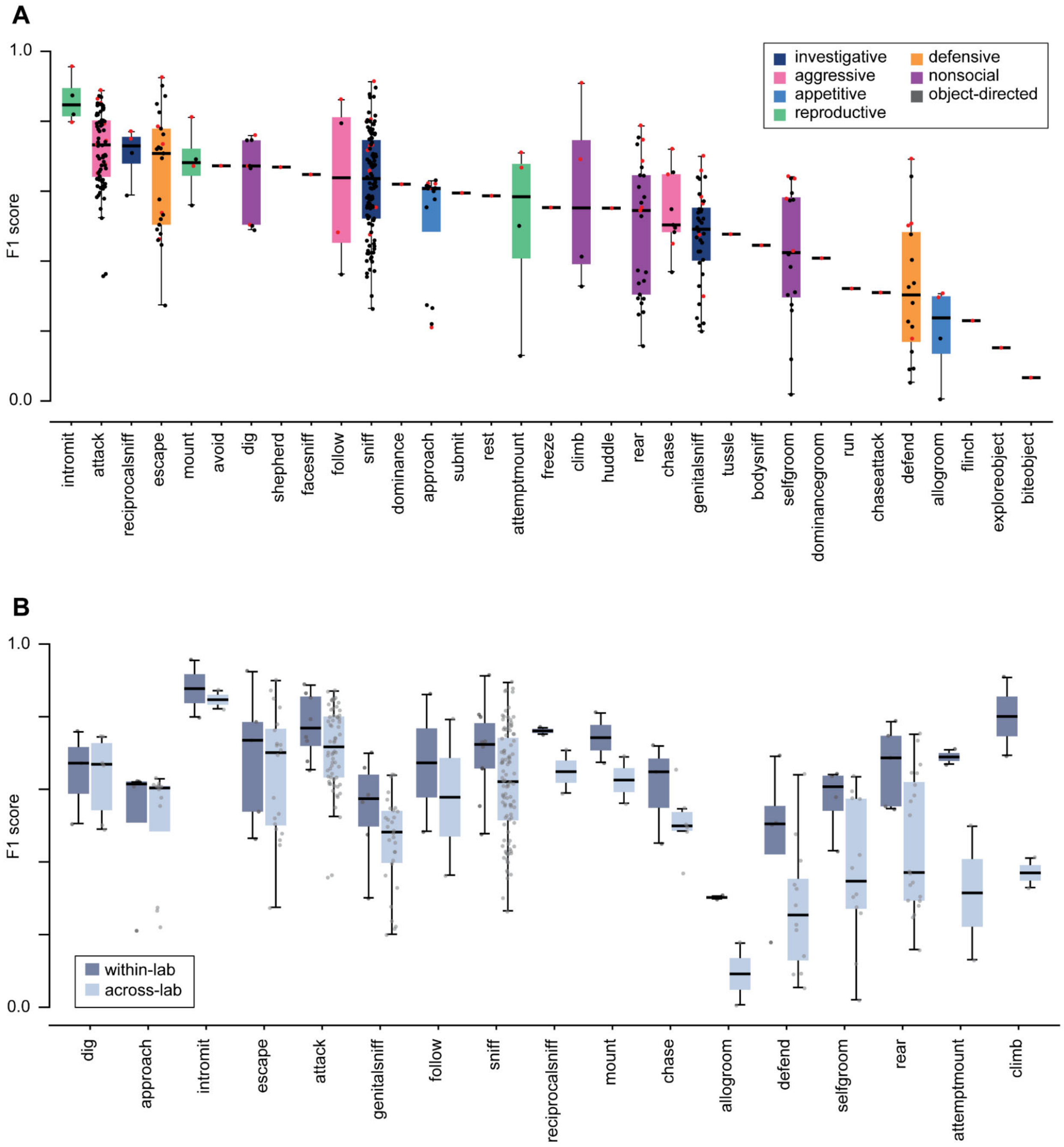

**Supplementary Figure 6. Within- and across-lab F1 scores. A)** Behaviors in the dataset sorted by median F1 score. Markers indicate within-lab (red) vs between-lab (black) f1 scores. **B)** Within- vs between-lab F1 scores for each behavior annotated by multiple labs, sorted by the within/between-lab F1 score gap. While some classifiers achieve comparable within- and cross-lab performance, others show a clear drop in F1 score. A large drop in cross-lab performance could reflect either lab-to-lab differences in how animals perform the annotated behavior, or lab-to-lab differences in how behaviors are operationally defined.

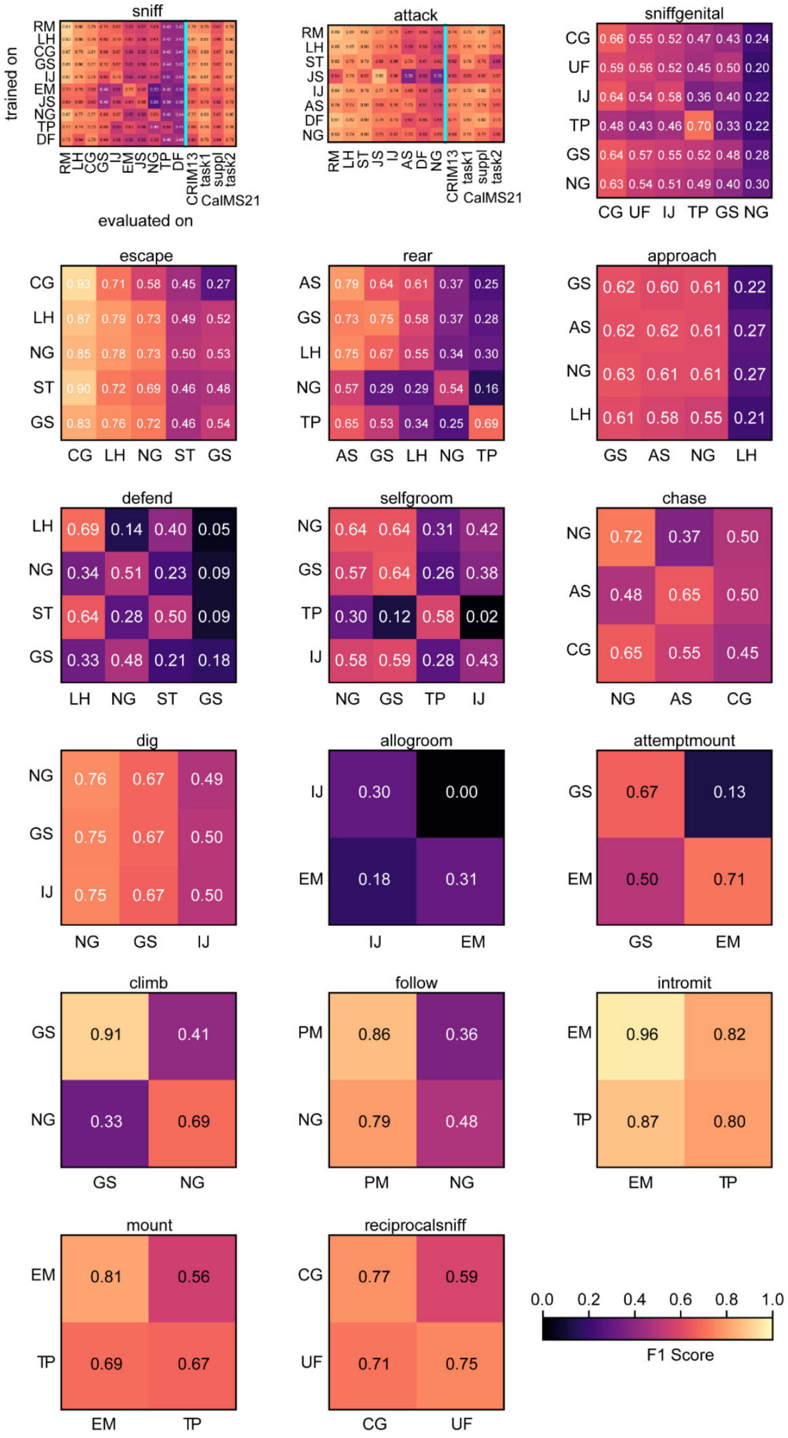

**Supplementary Figure 7. F1 scores for behaviors annotated by multiple labs.** For each behavior annotated by more than one lab, we compute the F1 score of each lab that trained a classifier for that behavior, when evaluating on the test set from each other lab that annotated that behavior. Each classifier's detection threshold (a single scalar value) was tuned on the training set for the evaluation lab.

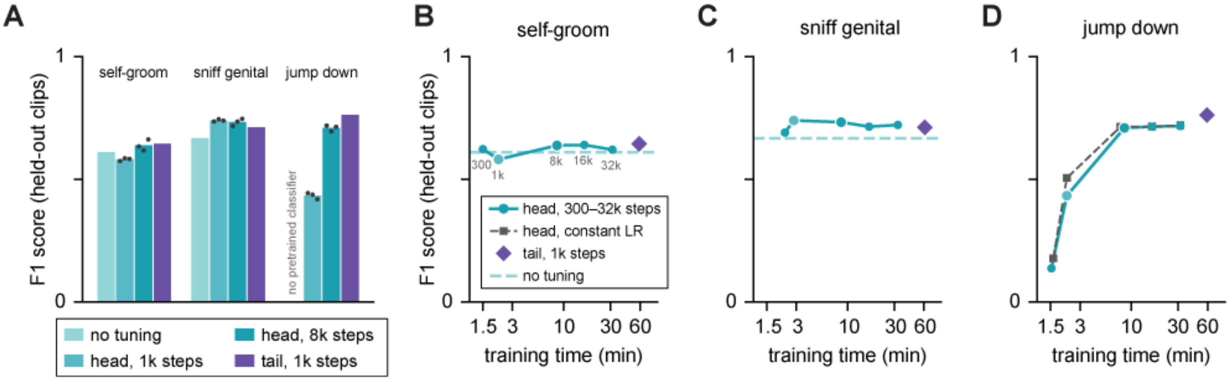

**Supplementary Figure 8. Accuracy and training cost of fine-tuning strategies. A)** F1 score of HiDRA classifiers on held-out lab's data, out-of-the-box ("no tuning"), after fine-tuning the output head for 1,000 or 8,000 steps ("head"; 258 parameters), and after fine-tuning the temporal filtering layers and output head for 1,000 steps ("head + trunk"; 4.8M parameters). All fine-tuning used annotations from two separate clips. F1 is computed on the four test clips concatenated, with each clip's decision threshold set on the other three; dots show individual training seeds. **B-D)** F1 score as a function of training time for self-groom **(B)**, sniff genital **(C)** and jump down **(D)**, fine-tuning the output head for 300 to 32,000 steps (step counts labeled in **B**); dashed lines indicate performance with no tuning, and the dark dashed line in **D** shows head fine-tuning with a constant rather than decaying learning rate. Head values are means of 3 seeds, each trained for its full step count; head + trunk values are single runs. Training time is wall-clock time to train the five-model ensemble on one GPU. Jump down has no pretrained classifier; the closest pretrained class (climb) scores near chance (F1 0.07) and is not shown.

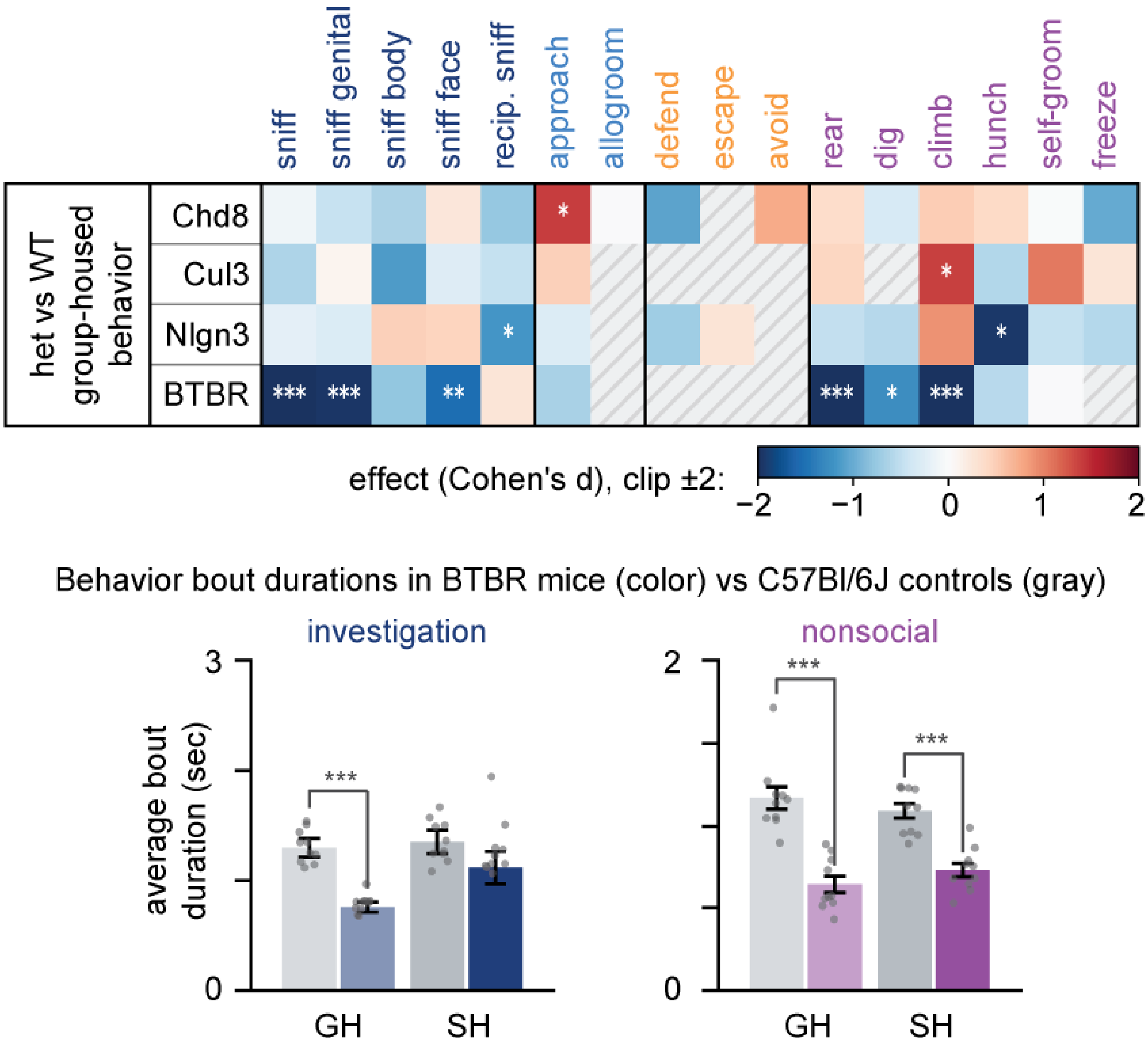

**Supplementary Figure 9. Interaction bout durations in ASD line study.** Above: difference in bout duration of behavior in heterozygotes compared to wild-type littermates (BTBR compared instead to C57Bl/6J) in the group-housed condition during the resident-intruder assay. Behaviors for which fewer than 3 bouts were observed in either experimental group were excluded from analysis. Color indicates effect size (Cohen's d.) Below: average duration of investigation and nonsocial behavior bouts for BTBR mice showing a reduction in bout durations. Stars denote significance in two-sided t-test, no multiple comparison correction.
